# Importance of depth refugia for reef resilience and intervention on the Great Barrier Reef under future climate change

**DOI:** 10.64898/2026.01.22.700983

**Authors:** Benjamin T. Grier, Takuya Iwanaga, Pedro Ribeiro De Almeida, Chinenye J. Ani, Samuel A. Matthews

## Abstract

Coral reefs globally face unprecedented threats from climate change, with the Great Barrier Reef (GBR) experiencing cumulative stressors and increasingly severe declines in coral cover from thermal stress events. Understanding drivers behind reef resilience to climate impacts is critical for conservation planning and intervention strategies. A scenario-based population modelling approach was adopted with larval connectivity dynamics and environmental factors to assess coral reef resilience across the GBR using projected conditions under five Global Climate Models (GCMs). Projected coral cover was analysed for each reef’s ability to maintain positive carbonate production budgets under future conditions, using coral cover as a proxy. Larval connectivity patterns did not correlate with increases in maintenance of positive carbonate budgets. Instead, reef depth emerged as the primary predictor, with deeper reefs (>10m) benefitting from reduced thermal exposure. These findings suggest that depth is a tangible and pragmatic reef characteristic to consider in future intervention practices for coral reef restoration. These results have important implications for reef management, indicating that depth should be considered as a key variable in conservation planning to maximize coral survival under continuing climate change.

## Introduction

Climate change is impacting species survival, persistence and ecosystem functions on coral reefs, which are vulnerable to increasing magnitude and variation in environmental stressors such as ocean temperature (Foster & Attrill, 2021; Gattuso et al., 2014). Mass bleaching and associated mortality events driven by climate change can cause coral population declines (Bozec et al., 2025; Hughes et al., 2017, 2018), possibly leading to coral-macroalgae regime shifts (Arif et al., 2022; Graham et al., 2015). Climate impacts on coral reefs are decreasing the ability of coral reefs to provide ecological functions such as benthic habitat and nutrient cycling (Forster & Zhang, 2021; Williams & Graham, 2019) and services such as food, protection, tourism and Indigenous cultural value (Eddy et al., 2021; Stoeckl et al., 2021).

Here, coral reef ecosystem resilience - the capacity to maintain coral populations and avoid ecosystem regime shifts - is conceptualised to depend on three interconnected mechanisms. First, the ability of individual corals to survive thermal stress. Second, the connectivity networks that enable recolonisation after disturbances (Hock & Mumby, 2017; Mumby & Hastings, 2008; Torres et al., 2024). Third, the maintenance, and accretion, of the carbonate framework that provides reef structure (Perry et al., 2012). This investigation aims to assess how these mechanisms may interact under projected climate conditions to identify the characteristics of resilient reefs in the Great Barrier Reef (GBR) ecosystem.

The GBR is the world’s largest coral reef ecosystem consisting of thousands of individual reefs interconnected through larval dispersal pathways mediated by hydrodynamic, spatial, and ecological processes (Cappo & Kelley, 2000). The GBR faces severe projected coral cover declines under climate change (Bozec et al., 2022, 2025; Condie et al., 2021). Coral cover declines will be compounded by ocean acidification, tropical cyclones, and crown-of-thorns starfish outbreaks (De’ath et al., 2012; Fox-Kemper et al., 2021; Seneviratne et al., 2021; Sweatman et al., 2011; Wolff & Anthony, 2018). The 2023-24 Summer mass bleaching event may be signalling a shift towards this reef future, with some level of bleaching recorded on 79% of surveyed reefs (total 1001 surveyed reefs), representing the most widespread bleaching event on the GBR and largest single year decline in the history of the Long Term Monitoring Program (LTMP, Australian Institute of Marine Science, 2024; Cantin et al., 2024).

At the individual coral scale, survival during thermal stress events depends on both environmental and biological factors that can either amplify or reduce the effects of thermal stress. Depth is a crucial factor that can provide a refuge from thermal stress (Álvarez-Noriega et al., 2018; Baird et al., 2018; Smith et al., 2014). Biological factors including coral size (Brandt, 2009), taxa (Álvarez-Noriega et al., 2018; McClanahan et al., 2005), and thermal tolerance through acclimation and inheritance (Humanes et al., 2022; Lachs et al., 2024) all influence survival outcomes. Corals demonstrate adaptive capacity to develop thermal tolerance that could delay widespread bleaching under moderate emission scenarios (Bairos-Novak et al., 2021; Humanes et al., 2022), representing a crucial mechanism for reef persistence.

Beyond individual survival, reef-scale resilience in part depends on connectivity networks that enable recolonization after disturbances (Angeler et al., 2023; Elmhirst & Hughes, 2009; Hock & Mumby, 2017). Larval source populations that have disproportionately numerous and/or strong connections to other reefs are essential for system-wide recovery, providing larvae that enable recovery at other reefs after disturbance (Hock & Mumby, 2017; Thomas et al., 2015). When disturbances impact major source populations, cascading effects reduce larval export and slow recovery throughout the connected network (Figueiredo et al., 2022; Hock & Mumby, 2017; Munday et al., 2009). Identifying and protecting source reefs has understandably been a major focus of research and management in recent years (GBRMPA, 2017).

Long-term reef viability requires not only coral survival and recolonization, but also maintenance of physical reef structures through positive carbonate budgets. Scleractinian corals build the primary habitat framework, but reef persistence depends on the balance between carbonate accumulation (Perry et al., 2015) and loss through bioerosion, physical damage (Kenyon et al., 2023), and dissolution (Eyre et al., 2014, 2018). This balance is influenced by coral cover and community complexity (Desbiens et al., 2021; Perry et al., 2012), environmental processes, and ocean chemistry (Eyre et al., 2018; Perry et al., 2015).

Recent advances in whole-ecosystem larval connectivity modelling (Choukroun et al., 2025) and coral thermal response research (Bairos-Novak et al., 2021) enable integrated investigation of how multiple factors interact to determine climate resilience. By incorporating the latest coral population models (Ribeiro de Almeida et al., 2024), ecosystem-wide connectivity data, and updated climate projections (Dixon et al., 2022), we can examine which factors most strongly influence reef persistence under different scenarios. While connectivity has been emphasized as critical for GBR resilience (Hock & Mumby, 2017), and remains an important consideration, our integrated modelling approach reveals that depth emerges as a dominant predictor for maintenance of reef coral populations and positive carbonate budgets under current projections of climate change.

## Methods

### Reef definitions

The spatial extent of reefs represented in this investigation follows reef definitions available via the Great Barrier Reef Marine Park Authority (GBRMPA). Reef definitions outline 3806 reefs across the GBR (Figure 1; GBRMPA (2022)). Prior analyses by Bozec et al. (2022) provided estimates of coral carrying capacity for each reef that were adopted by this study. Depths for each reef were extracted using 10m resolution bathymetry data made available in GBRMPA (2020). For each reef, bathymetry data were spatially matched with the 3806 reef outlines through geometric intersection. All depth values were converted to positive values (depths below sea level). The median depth was used to provide an indicative depth value for each reef. While the bathymetry data are said to be accurate up to 15m from Mean Sea Level (GBRMPA, 2020), there are many sources of uncertainty in the depth estimates used for this study. Notably, bathymetry data were derived from satellite imagery and can be limited in their alignment with the established GBRMPA reef outlines used to characterise reef locations and their areal extent. Further limitations are outlined in the Discussion. Code to extract depth estimates, and the resulting depth estimates themselves, are available at Iwanaga et al. (2025).

**Figure 1:**
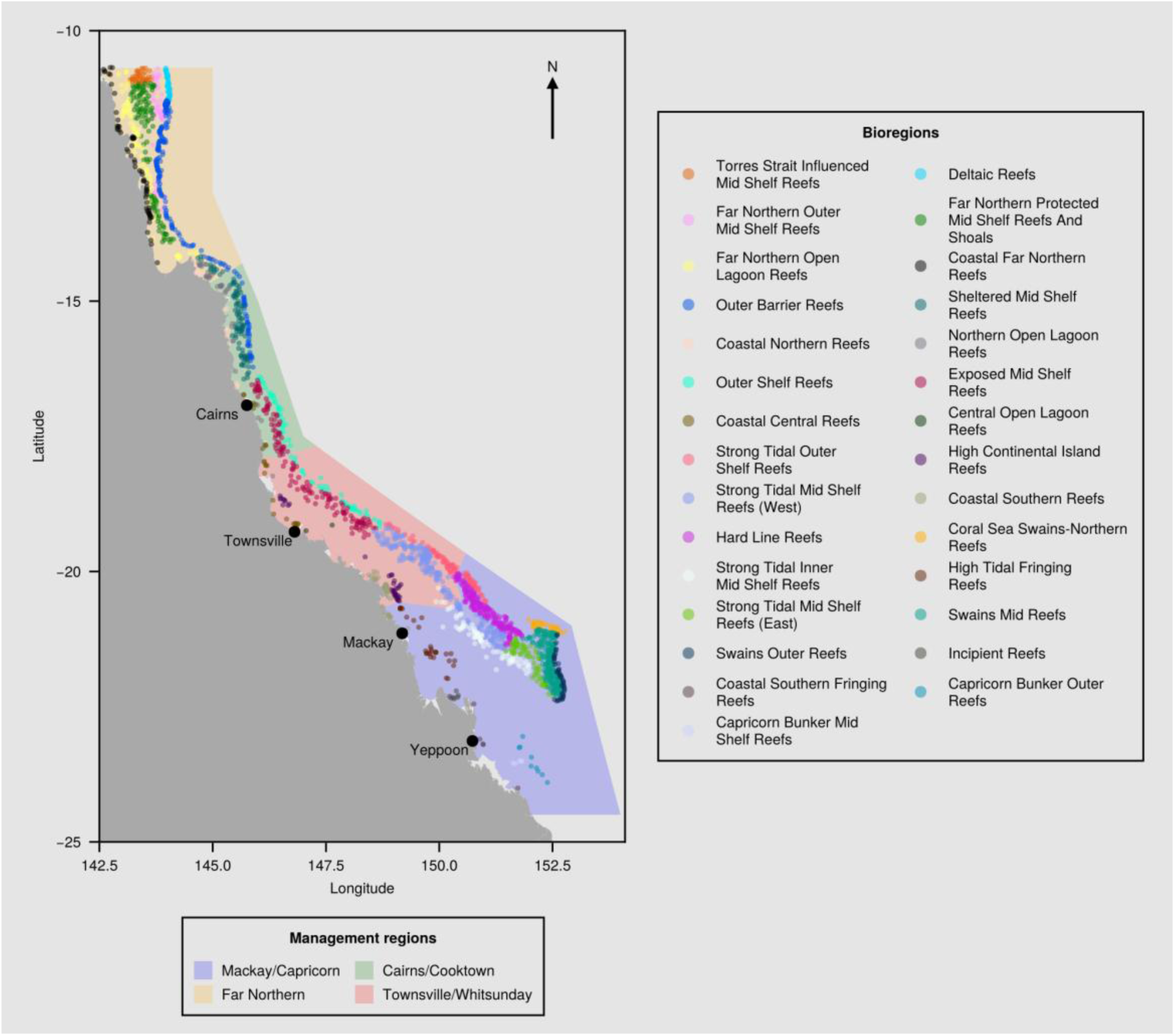
Map of the Great Barrier Reef region, including GBRMPA Management areas (polygons) and all considered reefs (points). Colour-coded polygons outline GBR marine park management regions: Far Northern (orange), Cairns-Cooktown (green), Townsville-Whitsunday (blue) and Mackay-Capricorn (red) (GBRMPA, 2007). Reef centroids are marked individually for visual clarity and are coloured by bioregions (GBRMPA, 2024). All 2197 reefs included in this investigation are represented in this figure. Annotations represent major coastal settlements along the GBR.

A Quality Assessment and Control process identified reefs with potential data limitations: reefs with no overlapping bathymetry data, reefs with minimum depth values above sea level (indicating potential data artifacts), and reefs where bathymetry pixels covered less than 5% of the reef polygon area. This process removed 842 reefs, or 22%, from the initial set of 3806 reefs. Furthermore, reefs that are not associated with GBRMPA Management areas (GBRMPA, 2007) or reefal bioregions (GBRMPA, 2024) were removed. Bioregions which contained fewer than three reefs after these processes were also excluded. As a result, 1377 reefs were excluded, leaving a total of 2429 remaining for time series clustering. To ensure effective data analysis, reefs with incoming or outgoing connectivity values below the 5th percentile were removed from post-hoc analyses, leaving 2197 reefs remaining for display in results (Figure 1).

### Degree Heating Week trajectories

Future projections of maximum yearly Degree Heating Weeks (DHWs) for each reef were estimated from Sea Surface Temperature (SST) data for the GBR. Statistically downscaled 1km^2^ resolution SST data for five Global Climate Models (GCMs) were obtained from the HighResCoralStress database (Dixon et al., 2022, 2023). These five GCMs were EC-Earth3-Veg, ACCESS-CM2, ACCESS-ESM1-5, NorESM2-MM and GFDL-CM4. The selected GCMs are a part of the Sixth Phase of the Coupled Model Intercomparison Project (CMIP6), spanning four ‘model development families’ as defined by Brunner et al., (2020). Shared Socio-economic Pathway (SSP) 2-4.5 was used to represent the plausible emission pathway. It represents a moderate emissions scenario with continued CO_2_ emissions comparable to present day until 2050, after which emissions gradually decline (Hausfather & Peters, 2020; IPCC 2023).

These GCMs were selected based on evaluation by Grose et al., (2023), indicating their ability to recreate historical atmospheric temperature and precipitation conditions in the Australasian region. They span a range of Equilibrium Climate Sensitivity (ECS) values within the Intergovernmental Panel on Climate Change (IPCC) assessed likely range. The IPCC defines ECS as the long-term warming caused by a doubling of carbon dioxide above its pre-industrial concentration (Forster & Zhang, 2021). While Grose et al. (2023) do not consider oceanic data, they provide a region-specific selection mechanism which is used as an indicator of general performance of GCMs for the region. An overview of the selected GCMs is provided in Table 1. Note here that Grose et al., (2023) assessed GFDL-ESM4, whereas Dixon et al., (2022, 2023) provide data based on GFDL-CM4. This substitution is assumed appropriate given they share the same fundamental representation of the physical climate system. The use of these GCMs therefore allows for a range of possible futures to be explored, capturing uncertainty in future responses to CO_2_ increases (Figure 2; SFigure 1; (Grose et al., 2023; Shepherd et al., 2018)) and enables the assessment of modelled coral reef behaviour over a broad range of independently derived climate futures (Abramowitz et al., 2019; Brunner et al., 2020).

**Figure 2:**
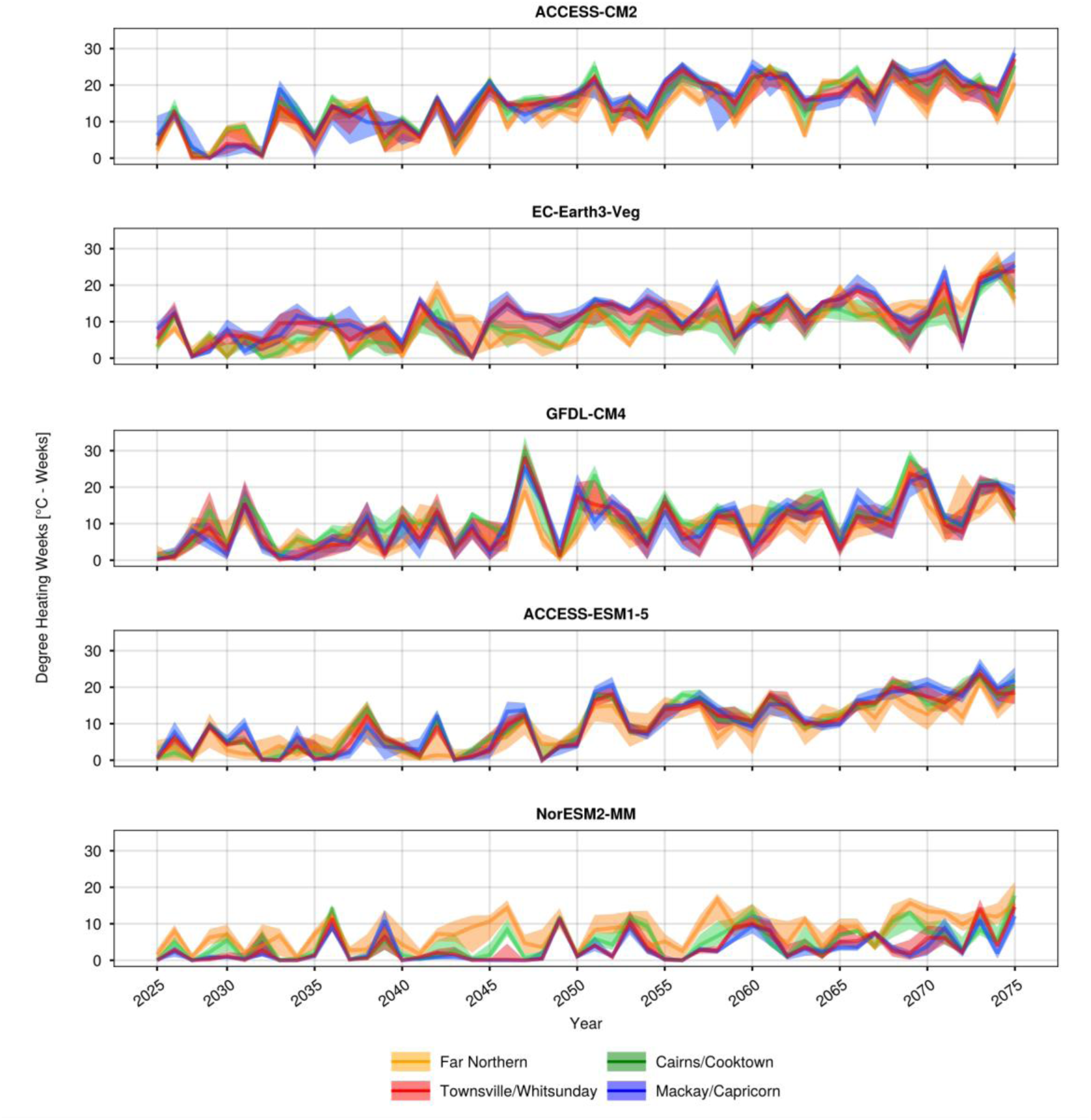
Degree Heating Week time series (in °C - Weeks) for the GCMs selected. Colours represent the different Management areas across the Great Barrier Reef: Far Northern (orange), Cairns-Cooktown (green), Townsville-Whitsunday (blue) and Mackay-Capricorn (red). Solid lines represent management-area median time series and bands represent 95th percentile confidence intervals.

**Table 1:**
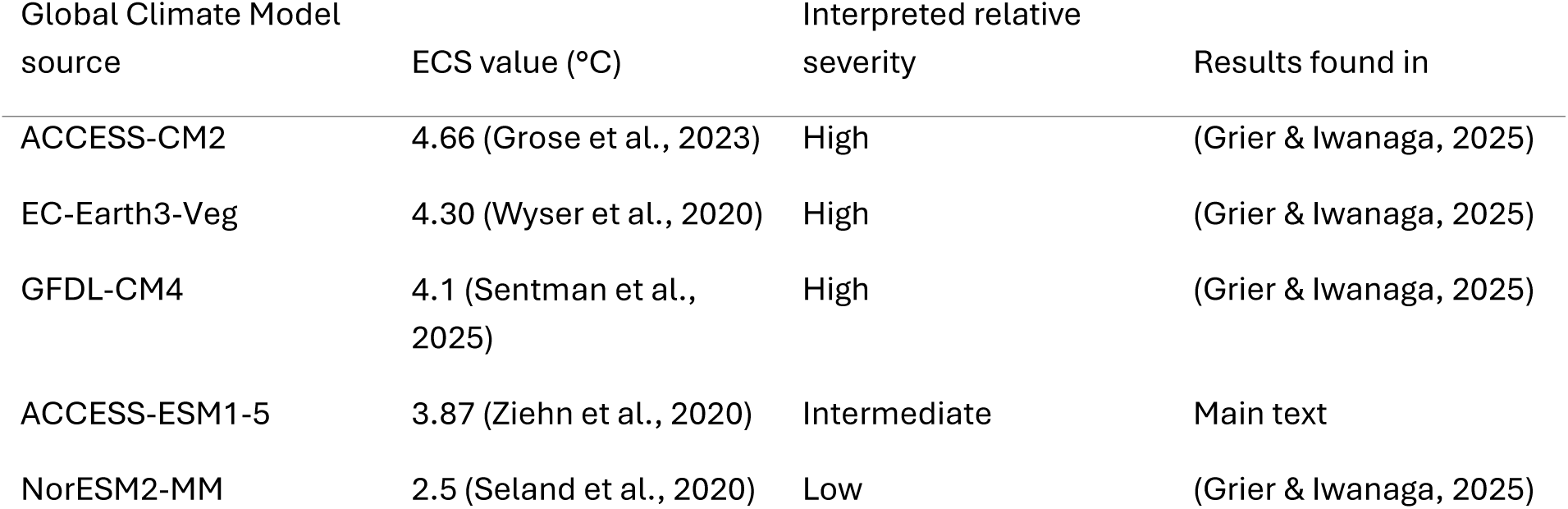
Information for each GCM used in this investigation including Equilibrium Climate Sensitivity (ECS) value, interpreted relative severity and where results for an individual GCM results are found. GCMs in context with assessed plausible ranges of ECS values are displayed in (SFigure 1). ACCESS-ESM1-5 has an ECS closest to the midpoint of the likely ECS range and was taken as an intermediate model with results displayed in the main text.

Daily SST data (Dixon et al., 2023) were used to calculate climatological baselines and “HotSpot Anomaly” values for each pixel following NOAA methodology (G. Liu et al., 2017; Skirving et al., 2020). Additional information about DHW calculation methods is provided in the supplemental material. DHW values on a per-pixel basis at 1km^2^ resolution were calculated as the 12-week rolling sum of HotSpot SST Anomaly values that are greater than 1°C, expressed in units of °C-weeks. The 12-week accumulation period reflects the cumulative nature of coral thermal stress. Brief temperature spikes may be tolerable, but sustained or repeated exposure to elevated temperatures causes progressively greater physiological stress and bleaching severity. DHW values integrate both the magnitude and duration of thermal stress: a moderate anomaly (e.g., 1°C) sustained for 12 weeks produces the same DHW value as a severe anomaly (e.g., 3°C) lasting 4 weeks, though the biological impacts may differ. DHW estimates for each indicative reef (see GBRMPA (2022)) were then determined by taking the mean DHW value for all pixels touching each reef outline. The process described here was conducted for all 2197 reefs to create DHW projections spanning from 2025 to 2075.

### Assessment framework

An exploratory scenario modelling approach was adopted for this investigation. The method involves exploring a wide range of potential futures and model parameterisations under different assumptions of climatic and ecological conditions. Exploratory modelling approaches have gained favour in recent years to explore the implications of deep uncertainty; where there is little or no agreement on the key drivers of a system, the extent of the relationship, how best to represent them or their probability distributions (Conradt et al., 2024; Lempert et al., 2013). The approach embraces uncertainty as an inherent characteristic of complex environmental systems (Lempert et al., 2013; Lempert & Groves, 2010). Exploratory scenario modelling is a core methodology underpinning the development of robust and adaptive management strategies (Haasnoot et al., 2013).

Recent critiques have raised concerns around the use of calibrated ecological models - which often have poor predictive capacity - for decision support (Lubiana Botelho et al., 2025). Here, the focus is on exploring the climatic and physical context that pertain to management decisions to understand the range of conditions under which conclusions hold true, rather than providing precise forecasts of future state. These environmental factors often set the context which influences ecological outcomes and have more manageable uncertainties than the precise ecological parameterisations (Reichert & Borsuk, 2005).

All data preparation, modelling and analyses were performed using the Julia programming language (Bezanson et al., 2017), with visualisations created using the Makie plotting ecosystem (Danisch & Krumbiegel, 2021). Code used to process DHW data, perform analyses, and create figures for this investigation are openly published and can be found in Grier and Iwanaga (2025).

### Connectivity modelling

The coral larval connectivity model used in this study comprises 3D hourly velocities from the 1 km grid resolution hydrodynamic model (GBR-1 version 2.0, Steven et al., 2019), as input to OceanParcels (Delandmeter & Sebille, 2019), a particle tracking model. The connectivity model was used to simulate larval dispersal and settlement of broadcast-spawning Acropora corals in the GBR, following annual mass spawning events in 2015, 2016, 2018, 2019 and split-spawning events in 2017 and 2020. Split spawning occurs when the annual reproductive event of a coral population is split over two consecutive months. Coral mass spawning usually occurs 4-6 days after the full moon during the spring-summer transition in the GBR with variations across species and locations. The spawning days identified are: 2015 - November 30, December 1 and 2; 2016 - November 18, 19 and 20; 2017 - November 8, 9 and 10, December 8, 9 and 10; 2018 – November 18, 19, 20; 2019 – November 16, 17, 18; 2020 – November 5, 6 and 7, December 4, 5, 6.

For each spawning night, particles were released from random locations within each reef polygon as neutrally buoyant virtual larvae every 27 seconds from 8:00 pm to 11:00 pm at 2.25 m below the surface. In each of the reef polygons, 401 particles were released resulting in a total of 1,526,206 particles every night of spawning across the GBR. The model was run for 30 days with a 3-minute timestep, and the locations of the released particles were updated every 15 minutes until the end of the simulation. Larvae were assumed to become competent to settle from 4 to 28 days post release with a daily mortality rate of 0.4 (Ani et al., 2024), which was implemented in 15-minute timestep as probabilistic mortality, corresponding to 0.0053 mortality rate per timestep. To incorporate the stochastic sensory behaviour of larvae during settlement, a 0.01 probability of settlement is assumed when larvae are over a reef during the competency period per 15-minute timestep. The information obtained on larvae settlement was used to compute a connectivity matrix among reef polygons for each spawning night (see Ani et al. (2024) for more details). A total of 24 connectivity matrices were generated for the spawning nights that were considered.

Larval transition matrices indicate the probability of larval transfer between pairs of reefs in the GBR, herein referred to as connectivity strengths. Indicative connectivity strengths are implemented in ADRIA-CoralBlox (described in the next section) as the mean of matrices for each location (i.e., mean of all 24 spawning days). While the connectivity strengths do not vary temporally within each ADRIA-CoralBlox simulation, spatial variability enables investigations into the influence of connectivity strength and its role in driving recovery processes.

To assess the impact of reef connectivity on modelled outcomes, connectivity was summarised by incoming connectivity (weighted incoming connectivity), outgoing connectivity (outgoing connection strength), and larval retention probability for each reef. Weighted incoming connectivity was calculated by summing the product of each connection strength incoming to a target reef (excluding retained larvae from the target reef) and the mean timeseries absolute coral cover (km^2^) of each matching source reef. This metric provides an approximation of overall larval flow into a reef, with high values indicating the target reef is supplied high volumes of larvae via strong connections and low values indicating the target reef has weak (or no) incoming connections or is supplied by small source populations. A limitation of weighted incoming connectivity is that it does not incorporate the proportion of the source reef’s population that is reproducing and producing larvae, instead it takes total coral cover area as an indicator of larval productivity. Outgoing connection strength was calculated as the sum of all outgoing transition probabilities from a target reef. Outgoing connection strength has a theoretical maximum of one (all larvae reach settlement on sink reefs), however the large size of the GBR combined with larval mortality result in much lower outgoing connection strength values. Reef larval retention probability indicates the proportion of spawned larvae that settle on their natal reefs.

### Scenario modelling

Two modelling systems were employed to explore scenarios: ADRIA (the Adaptive Dynamic Reef Intervention Algorithms platform) and CoralBlox, referred to as the coupled framework ADRIA-CoralBlox. ADRIA is a multi-criteria decision support platform designed for the assessment and analysis of coral reef intervention strategies under uncertainty (T. Iwanaga et al., 2023). The ADRIA platform provides functionality for scenario generation and analysis as well as providing technical infrastructure to couple environmental and ecosystem representations. CoralBlox is a computationally efficient coral population model. It simulates functional/morphological groups of corals over their life cycles, competing for available space over time (Ribeiro de Almeida et al., 2024).

The coral demographic representation and associated model implementation are further detailed in the supplementary material. Of note is that the representation of bleaching mortalities followed Bozec et al., (2022) and Baird et al. (2018) which applies a scaling coefficient to incorporate the amelioration effect that depth may have. This depth-scaling reduces the proportion of coral colonies impacted by high heat stress as depth increases. The impacts of tropical cyclones, wave activity, Crown of Thorns Starfish or coral disease were not included in this investigation in order to focus on assessing reef resilience under possible future warming scenarios.

ADRIA-CoralBlox was run for 4096 scenarios for each GCM (20,480 total model runs). Here, a scenario represents a unique combination of assumed environmental (namely one of the five DHW trajectories) and ecological factors including coral growth and mortality rates, and coral thermal tolerances. Scenarios were generated using the low-discrepancy Sobol’ sampling method (Sobol’, 1967, 1993) with Owen Scrambling (Owen, 1995), a randomised Quasi Monte Carlo approach which balances scenario exploration with improved convergence properties compared to pseudo-random sampling. A detailed comparative analysis of sampling methods can be found in (Kucherenko et al., 2015; L’Ecuyer, 2018). For scenario assessment, coral cover time series for each model run were scaled to represent cover as a percentage of the total estimated reef area. The scenario-median time series for each reef was used for further analyses. Time series of coral cover were then clustered based on their behaviour. Assessment metrics used in analyses can be found in STable 1.

### Time series clustering

Reefs were clustered based on the behaviour of their median coral cover time series. Put simply, reefs that have similar trends and trajectories are more likely to be clustered together. Reefs are clustered into Low, Medium, or High, indicating their relative ability to maintain coral cover over time. Reefs in each time series cluster were compared to identify characteristics that differ between Low, Medium and High clusters. These characteristics include reef median depth, mean time series DHW, weighted incoming connectivity, outgoing connection strength, larval retention probability and reef carrying capacity (coral habitable area).

The Complexity-Invariant Distance (CID) between each time series was calculated to identify similarities in temporal behaviour (Batista et al., 2014). The CID applies a correction factor to the Euclidean distance between pairs of time series to obtain a measure of their similarity. The adopted approach ensures reefs are clustered based on their temporal dynamics as opposed to, for example, naively clustering by their mean coral cover. Time series clustering is achieved using k-medoids clustering (cf. Steinmann et al., 2020). Further details on the time series clustering approach are found in the supplementary materials.

Time series clustering was applied over the time frame between 2025 and 2075. The clustering was conducted at three spatial scales: (1) the whole of GBR; (2) each GBR management area; and (3) for each reefal bioregion (defined in GBRMPA (2024)). Analyses were conducted at the different spatial scales as, conceptually, the temporal behaviour constituting “low”, “medium” or “high” coral cover may differ at different spatial scales. Analyses at the whole-of-GBR scale were found to lead to identical conclusions regarding reef characteristics which influence coral cover and so for brevity are not shown here.

To compare time series clustering results across GCMs, management area scale clustering results are used to preserve coarse scale latitudinal differences between regions, while maintaining ease of comparison and visualisation. These comparisons include quantification of differences between low and high cover clusters across management regions under each GCM using the CID between cluster median time series. To assess how different reef characteristics influence time series clustering at a finer spatial scale, bioregion results are used to capture patterns occurring within individual groups of reefs. Results of all analyses at all spatial scales can be found at Grier and Iwanaga (2025) for those interested.

### Partial Dependence Analysis

Partial Dependence Analysis (PDA) was conducted to understand relationships between features and bioregion-scale time series cluster predictions using a Random Forest (RF) classifier (Friedman, 2001). The analysis utilised the reef characteristics of interest as the predictors, including the estimated median reef depth, mean DHW for the given GCM, log10-weighted incoming connectivity, outgoing connectivity, and log10-transformed outgoing-to-incoming connectivity ratio. Conceptually, PDA is related to a Regional Sensitivity Analysis (Hornberger & Spear, 1981; Pianosi et al., 2016), as model results are assessed to map the influence of 𝑋 - the predictors or model factors - on 𝑦 - the modelled outcomes according to a metric of interest.

The full dataset consisted of the time series cluster assignment for each of the 2197 reefs under the each GCM (totalling 10,985 samples). The dataset was randomly split into training (60%) and testing (40%) subsets to train and assess classification performance. The RF classifier achieved 80% accuracy on the training set and 60% on the test set. A 3-fold cross-validation was also conducted, arriving at a similar 59% accuracy rate. F1 performance scores for Low, Medium, and High clusters were determined to be 0.70, 0.42 and 0.63 respectively, with precision of 0.63, 0.48 and 0.64 when evaluated with the test set (consisting of 4394 samples) (STable 3; STable 4). Lower performance with the Medium cluster reflects the inherent difficulty of classifying transitional states in natural systems. The consistent level of accuracy across test and cross-validation demonstrates the generalised performance of the RF classifier, while not accurate, is still sufficiently informative for the purpose of assessing the influence of reef characteristics.

Partial dependence analysis was performed by applying the trained model to the test dataset. Values for individual predictors were systematically varied (one-at-a-time) across a grid of 50 equally spaced points spanning the predictor’s range, while all other features were left at their modelled values. The procedure creates (1) a modified test dataset, where the target predictor was set to the grid value for all samples, (2) probabilistic predictions for all three cluster classes using the trained RF classifier, and (3) the mean predicted probability for each class across all test observations. The process isolates the marginal effect of each predictor by averaging over the natural distribution of all other representative reef characteristics in the test dataset. Additionally, to assess possible interaction between reef depth and connectivity metrics, two-way PDA was performed on grids along the ranges of depth and weighted incoming connectivity, and depth and outgoing connection strength.

### Carbonate budget analysis

Using coral cover time series for each reef, the number of years each reef maintains coral cover above a given percent threshold were calculated. The threshold represents the level of live coral cover necessary to maintain a positive carbonate budget (denoted here as 𝜃_0_to indicate the budget equilibrium). A greater number of years above 𝜃_0_suggests greater resilience and opportunity for recovery.

Recent analyses suggest a general threshold of ∼17% live coral cover in the case of the mid-shelf and offshore GBR reefs (Desbiens et al., 2021). Given in-situ carbonate budgets are influenced by (1) a range of processes; (2) the variety of approaches that could be used to measure those processes; and (3) the uncertainty surrounding these, it would be unreasonable to assume all reefs adhere to a single precise 𝜃_0_value. Concurrently, it is difficult to ascertain a suitable 𝜃_0_value for each individual reef.

Given the uncertainties present, the proportion of modelled reefs that display capacity to maintain coral cover above a given 𝜃_0_ threshold is examined. An indicative range of 10 to 20% is used (inclusive, with a step size of 1%). The range was informed by Perry et al., (2015) and Desbians et al., (2021) providing plausible indications of the lower and upper bounds respectively. Robustness of the conclusions drawn to the uncertainty in the precise value of 𝜃_0_for all reefs can then be assessed by using the indicative range.

The relationships between the number of years above 𝜃_0_ and characteristics such as median reef depth and log10 total connectivity strength were assessed with Spearman’s rank correlation. Spearman’s correlation was adopted as distributions of reef depths and number of years above 𝜃_0_were expected to be non-normal and highly skewed. Furthermore, reefs are naturally of different types and experience different conditions, and so the monotonic trends (whether the relationship consistently increase or decrease) rather than the exact quantitative relationship was of interest.

## Results

### Reef trajectories across futures

There were substantial differences in the maintenance of coral cover post 2030 between the GCMs (Figure 3). Of the GCMs explored, NorESM2-MM had the lowest ECS (2.5°C) and reefs exhibit continuing recovery and decline dynamics up to 2075 under this hypothetical climate future. Unsurprisingly, recovery phases were predominantly seen in the reefs in the High cover cluster, with Medium and Low exhibiting lesser probability of such periods of recovery. The High cover cluster exhibits greater resilience despite encompassing a broad range of initial coral cover conditions at 2025 (Figure 3). Generally, recovery and increases in coral cover were visible in simulated results up to 2045. Under NorESM2-MM, the Far Northern Management area was found to have the most similar temporal behaviour between Low and High clusters compared to other Management areas (STable 2) due to the low degree of coral cover overall (Figure 3).

**Figure 3:**
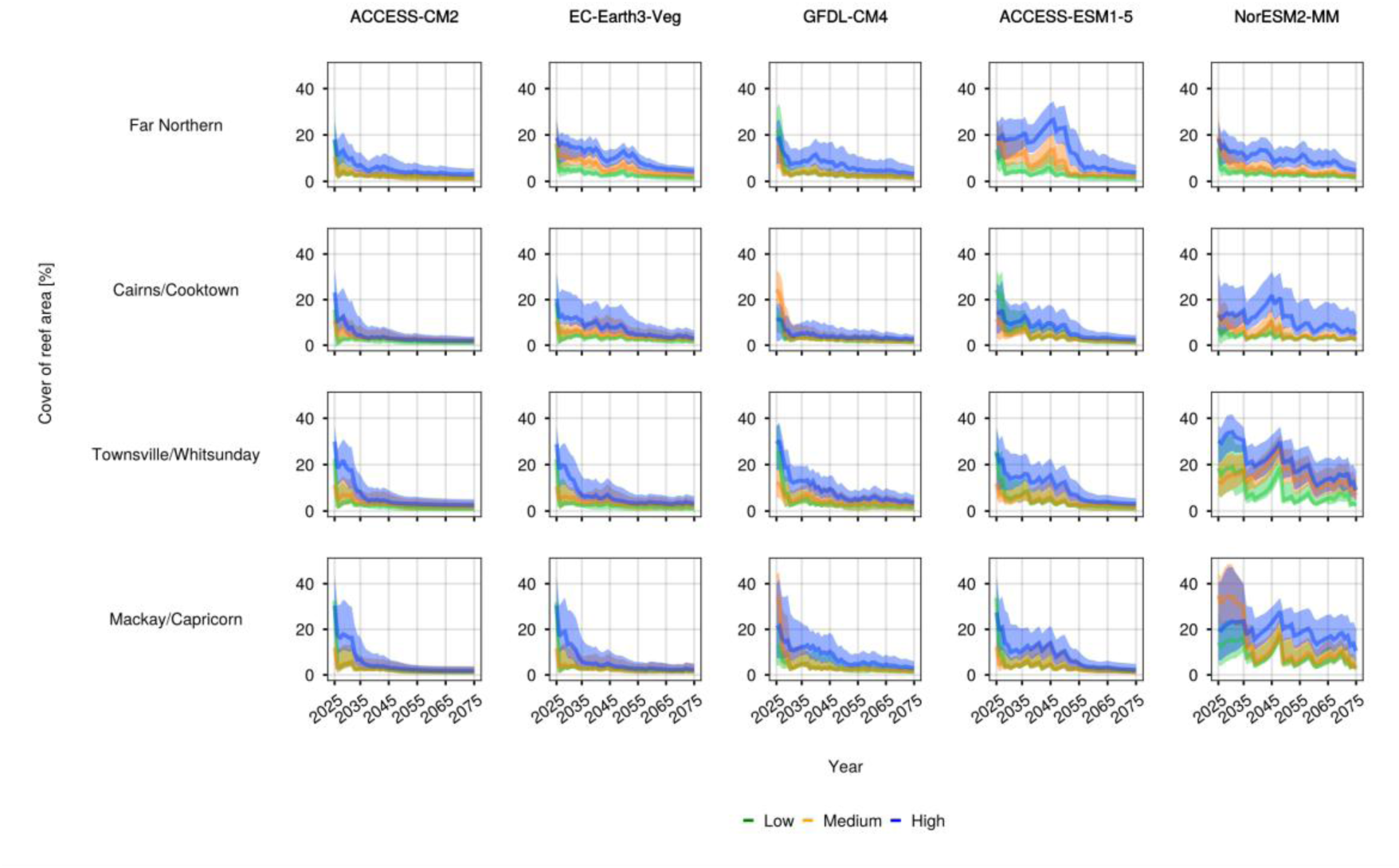
Time series of coral cover (relative to reef area) clustered within each Management area (rows), for each GCM analysed (columns) based on temporal behaviour over 2030 and 2060. Time series were clustered into “Low” (green), “Medium” (orange), and “High” (blue) clusters indicating their relative median coral cover. Solid lines represent cluster median time series and bands represent 95th percentile confidence intervals.

Intermediate levels of coral cover maintenance can be seen in ACCESS-ESM1-5 (ECS value of 3.87°C) when compared to the other GCMs (of higher and lower ECS values). Under these intermediate conditions (in relative terms), there is potential for reefs in the High cluster to maintain coral cover levels around 20% to 2050. The highest peak DHW levels occur under GFDL-CM4 between 2045 and 2050, while greater increases in overall DHWs time series occur under ACCESS-CM2 (Figure 2). Under conditions represented by high ECS value GCMs - specifically, ACCESS-CM2 (4.66°C), EC-Earth3-Veg (4.30°C) and GFDL-CM4 (4.1°C) - all reefs are largely in decline across all management areas from 2035 (STable 2).

Comparing across GCMs, it is common for reefs to switch between clusters, including from Low to High and vice versa (SFigure 5). Of 2197 reefs, 21% (462) of reefs have consistent cluster assignment across all GCMs. Of the 462 consistently assigned reefs, 23% are within the Far Northern Management area, with 1%, 55% and 21% occurring in Cairns-Cooktown, Townsville-Whitsunday and Mackay-Capricorn Management areas respectively (SFigure 4). Of the 462 consistently assigned reefs, 34%, 18% and 47% are clustered into Low, Medium and High clusters respectively. High levels of assignment variability across GCMs are expected due to spatial differences in each given GCM’s DHW trajectories.

### Bioregion scale clustering

Relationships between reef characteristics and bioregion-scale time series clusters were found to be similar across GCMs. Across bioregions, reefs in the High cluster not only had greater levels of coral recovery but displayed persistence to 2050 (SFigure 18). Qualitative representations of bioregion scale clustering results across assessed reef characteristics can be found in the supplementary material, with Partial Dependence Analysis results displayed in the main text.

### Partial dependence

Partial Dependence Analysis revealed reef median depth, carrying capacity and mean time series DHW level have the highest feature importance scores for bioregion-scale cluster assignment, accounting for 32%, 16% and 15% predictive importance respectively (Figure 4; Panel A). Outgoing connectivity strength, Source-cover weighted incoming connectivity and larval retention probability account for ∼14%, 14%, and 10% respectively. Reefs have the greatest probability of being clustered into Low if shallow (<10m), and High at greater depths, with medium cluster reefs exhibiting a lower, and more stable, assignment probability across depth ranges (Figure 4; Panel B). A large increase in the probability of a reef being categorised as High can be seen between 10 and 15 meters depth. Across mean DHW values, assignment probability is roughly equal across the three clusters up to 5 DHW. A sharp delineation can be seen > 5 DHW when comparing High with other clusters, with lower assignment probability (∼25%) compared to Medium and Low (∼40%) (Figure 4; Panel B).

**Figure 4:**
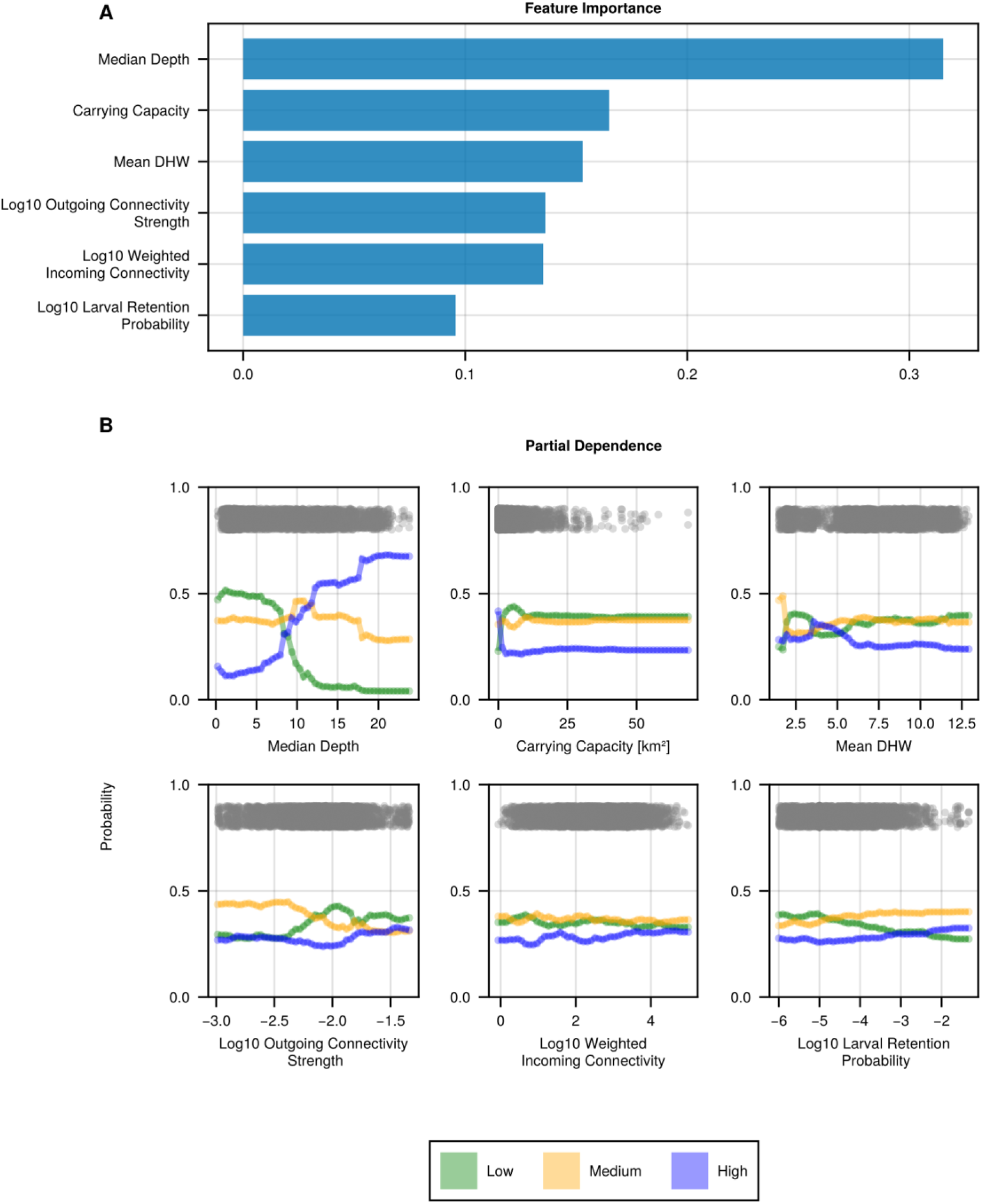
Feature importance scores (Panel A) of the Random Forest classifier, and Partial Dependence Plots (PDPs) for reef characteristics of interest indicating the probability of a reef being classified to one of the three cover clusters (Panel B). PDPs indicate the probability of a reef from the test dataset being clustered into each cover cluster depending on the predictor value on the x axis, when adjusted for values of other predictors. Green, orange and blue lines indicate cluster assignment probability for Low, Medium and High clusters respectively. Transparent grey points above probability lines represent predictor values present in the test dataset, providing a rough indication of predictor value distribution across assessed reefs.

First-order PDA showed minimal influence of incoming and outgoing connectivity metrics on cluster assignment probabilities. Two-way analyses revealed little interaction between depth and incoming and outgoing connectivity metrics for reef clustering probabilities (Figure 5; SFigure 6). Reefs with a median depth around 15m may benefit slightly from high incoming and outgoing connectivity, with higher probability of High cover cluster assignment compared to 15m - depth reefs with lower connectivity.

**Figure 5:**
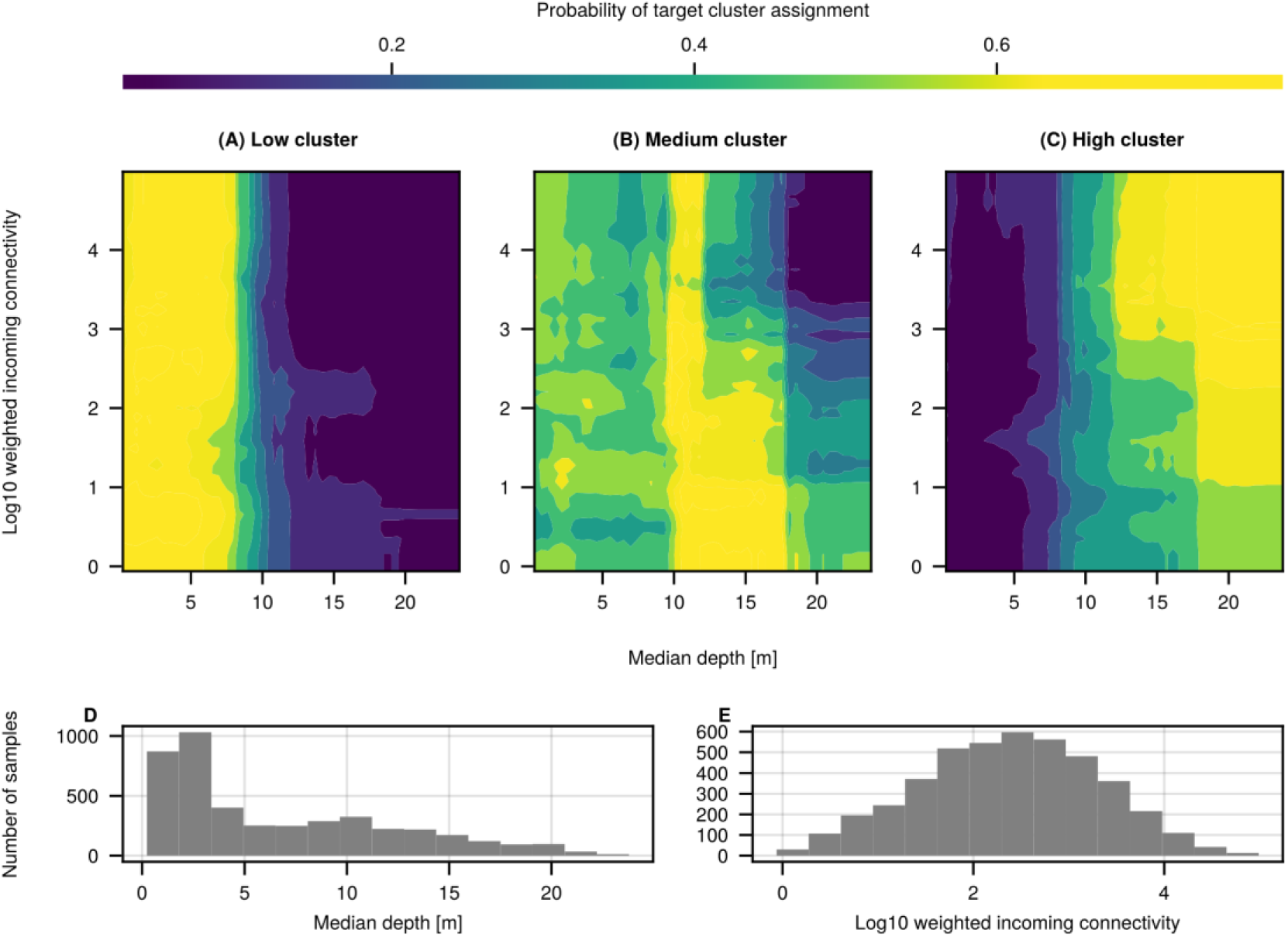
Partial Dependence Plot indicating the estimated probability (via Random Forest classification) of cluster assignment along interacting values of median reef depth (meters) and log10-weighted incoming connectivity. Colour (ranging from dark purple to yellow) indicates the probability of assignment to each cluster, with brighter shades indicating higher probability of assignment. Panels A, B, and C illustrates the assignment probability for Low, Medium, and High cover clusters respectively. The distribution of reef depth and incoming connectivity values are shown in Panels D and E.

Individual feature analysis for carrying capacity indicates that its influence on cluster assignment is concentrated at the low range of predictor values (Figure 4). This aligns with the right-skewed distribution of carrying capacity across reefs for these three predictor variables. Beyond this low-end range, partial dependence becomes near constant, indicating outlier values for these factors are not influential. Assessing random forest prediction accuracy across bioregions reveals increasing performance as mean bioregion reef depth increases (SFigure 7). Unexplained variation in the test dataset (33% of variation) may be concentrated in shallow bioregions where considered model predictors do not capture differences between clustered reefs.

### Reef carbonate budget

Reef median depth had a moderate positive correlation with the number of years a reef maintains coral cover above the assumed threshold required to sustain a positive carbonate budget (𝜃_0_). Greater median depth was a key characteristic defining reefs that sustain coral cover above 𝜃_0_ (Figure 6; SFigure 15; SFigure 16). Unsurprisingly, the number of years above 𝜃_0_substantially decreased as 𝜃_0_is increased, as does the Spearman’s correlation (SFigure 15). Decreasing correlation levels are likely due to an increasing number of reefs being unable to maintain coral cover above 𝜃_0_ as it is increased, with the distribution of years above threshold becoming increasingly right-skewed (Figure 6). In contrast, weighted incoming connectivity strength (log10 scale) was found to have no correlation with the number of years a reef maintains cover above the required thresholds (SFigure 15).

**Figure 6:**
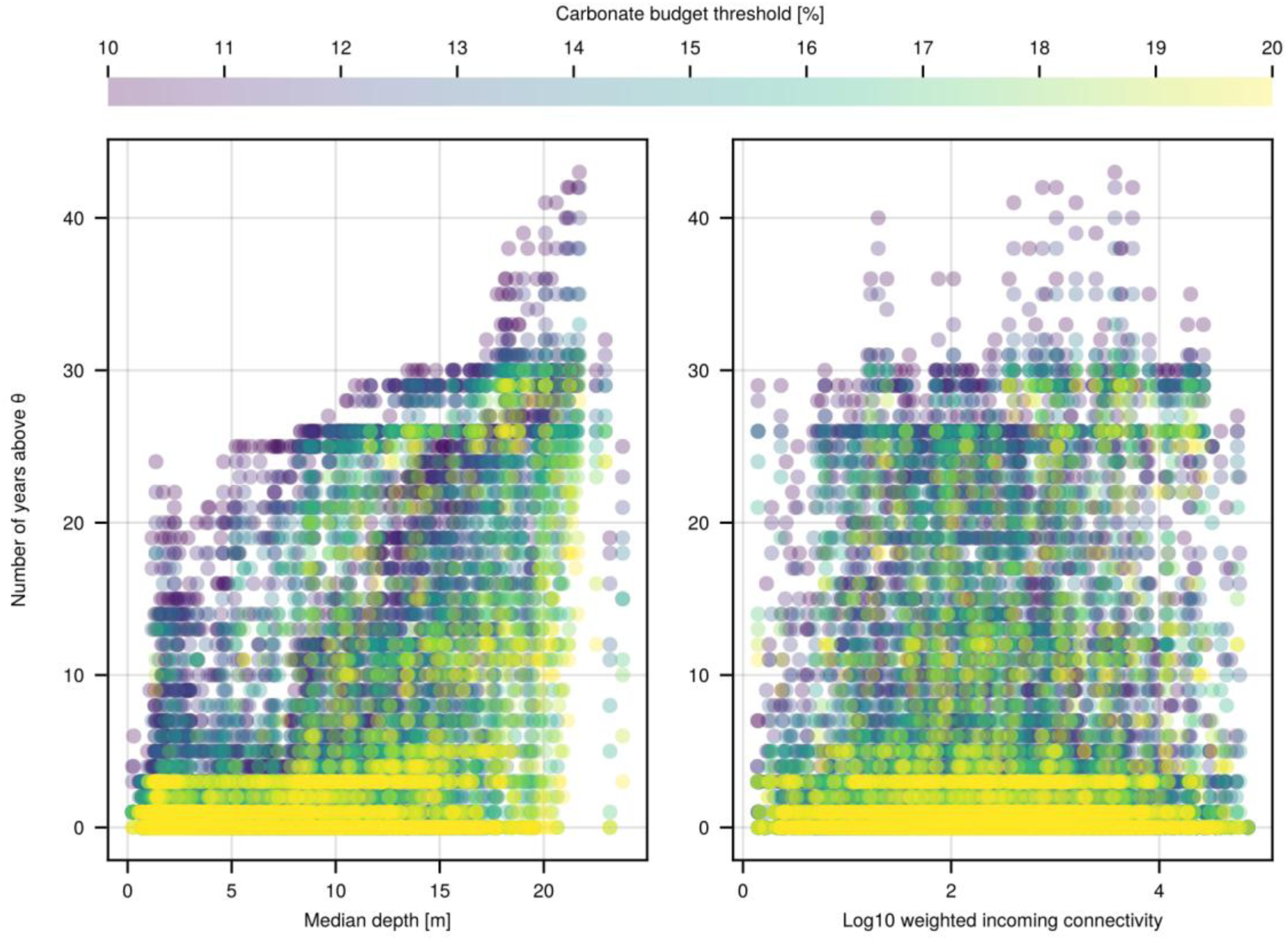
Number of years reef coral cover exceeds the assumed carbonate budget threshold in terms of coral cover (𝜃_0_) between 2025 to 2075 (y-axis) under ACCESS-ESM1-5. Left panel shows the relationship between reef median depth, while the right panel shows the same for log10-weighted incoming connectivity. Marker colour (purple to yellow) represents 𝜃_0_, with brighter shades indicating a higher 𝜃_0_. Deeper reefs have an increased number of years above the considered range of 𝜃_0_, while no relationship between weighted incoming connectivity and carbonate budget maintenance was found.

## Discussion

Results show increasing reef depth has a strong positive impact on modelled reef resilience in temporal behaviour and maintenance of positive carbonate budgets under possible future conditions. Reefs with a median depth of 10m or greater, with high incoming connectivity, are the most likely to be assigned to a high cover time series cluster. By itself, incoming connectivity alone does not correlate with maintenance of positive carbonate budgets. These findings highlight the importance of depth and incoming connectivity as coupled selection features when assessing potential reef resilience under future thermal stress.

Reefs in different temporal clusters largely do not differ in mean modelled DHW levels experienced and exhibit overall similar DHW time series. While reefs experience similar magnitude DHWs based on SST, the proportion of a reefs’ coral population impacted by heat stress events depends on reef depth. Results of this investigation suggest that for non-mesophotic reef ecosystems (i.e., < 30m), depth contributes greatly to reef resilience and maintenance positive carbonate budgets under future conditions.

Movement of shallow reefs towards negative carbonate budgets in the future may contribute to phase shifts in dominant sources of CaCO_3_ production. Reefs may switch away from coral-driven processes, and CaCO_3_ accretion, to encrusting algae driven CaCO_3_ production, or to bioerosion dominated systems with rapid erosion of physical reef frameworks (Dudgeon et al., 2010; Perry et al., 2008). Reefs at increased depths may be able to maintain coral driven CaCO_3_ production and accumulation into the future for longer.

Spatially these results indicate that regions of positive carbonate budget maintenance, under ACCESS-ESM1-5, and high depth levels were concentrated in the Far Northern, offshore Townsville/Whitsunday, and Mackay/Capricorn management areas (SFigure 9; SFigure 17). High cover reefs in the Far Northern management area were exposed to lower mean DHW levels than other reefs under ACCESS-ESM1-5 (SFigure 25), however similar spatial patterns were observed across GCMs for reefs clustered into the high cover cluster (SFigure 4).

These findings are not to be interpreted as accurate predictions of future resilience due to high uncertainty in input data, particularly depth and connectivity. While similar findings have been demonstrated in the Pacific (Baird et al., 2018; Smith et al., 2014) and Indo-Pacific (Bridge et al., 2014) regions, depth refugia analyses have often focused on the potential of mesophotic reefs (> 30m deep) as thermal refugia (Bongaerts et al., 2010; Holstein et al., 2015; Semmler et al., 2017). In contrast, Venegas et al. (2019) found no association between depth and the alleviation of heat stress across a range of shallow reefs in the Pacific.

This investigation supports the hypothesis that reefs at greater depths will exhibit characteristics of refugia and prolonged resilience (in terms of coral cover) under climate change compared to shallower reefs. Additionally, Liu, Callaghan and Baldock (2024) suggest that rubble instability is inversely correlated with reef depth, possibly limiting juvenile coral recruitment capacity of shallow reefs. A caveat to depth-resilience findings is the quality of the bathymetry data used to calculate median reef depth. As detailed previously, estimates of median reef depth are derived from satellite imagery (at 10m^2^ resolution) which can be subject to high degrees of uncertainty, both in terms of the estimated depth for each pixel as well as their alignment to the established GBRMPA reef outlines.

Current reef restoration and intervention efforts across the GBR, such as deployment of coral seeding, coral fragment devices and Crown of Thorns Starfish control, target reefs with sites in the 1-10m depth range (Howlett et al., 2021; Randall et al., 2021; Whitman et al., 2024). While there are no explicit justifications for this depth range, intervention depth is likely constrained by snorkelling and diving deployment and monitoring logistical accessibility, visual identification of sites from the surface, in combination with tourism return value. Of the assessed reefs occurring within the 1-10m depth range (1551 reefs), 59% (920) of reefs are consistently clustered into the Low and Medium cover clusters across the GCMs (at the GBRMPA management region scale), with many found to be unable to maintain positive carbonate budgets under projected conditions. PDA revealed a range of depths from 10-15m where the probability of a reef being clustered into the High cover cluster increased substantially compared to shallower depths. Considering this 10-15m depth range when assessing possible intervention sites may reduce the exposure of planted and seeded corals to high thermal stress, while still maintaining effective logistical access to sites.

It is a common conclusion in coral reef ecology that a primary facet of reef resilience is the supply of recruits that a reef receives following disturbance (Hock & Mumby, 2017; Mumby & Hastings, 2008; Torres et al., 2024). No direct, first order, relationship between weighted incoming connectivity and the ability of reefs to maintain positive carbonate budgets under future conditions was found. There has been contrasting evidence (Bode et al., 2018; Hock & Mumby, 2017) as to whether current connectivity modelling is able to identify reefs that are important for recovery processes, with findings dependent on the larval connectivity model used. These investigations were performed using prior generations of larval connectivity models parameterised for 2008-2013 (Hock & Mumby, 2017) and 1996-2002 (Bode et al., 2018) time periods, without considering the influence of reef depth on resilience. Using updated 1km^2^ resolution larval connectivity across the whole GBR, with a limited temporal sample of particle flows, it was found that larval connectivity alone may not be enough to support reef resilience and recovery but should be considered in combination with more influential factors such as depth.

Findings on the importance of connectivity likely differ from literature due to the highly uncertain nature of larval connectivity modelling (Choukroun et al., 2025; Vogt-Vincent et al., 2023) and the way larval flows and juvenile recruitment are implemented in ADRIA - CoralBlox. Additionally, and the high magnitude of early bleaching events (2025-2035) may increase the influence of temperature mediators (i.e. depth) in our simulations decrease the breeding capacity of strongly connected reefs, resulting in reduced importance of connectivity in recovery. Analysing daily connectivity data Vogt-Vincent, Mitarai and Johnson (2023) suggest that time-mean connectivity may not be suitable for capturing the seasonal - ecological time scale connectivity between locations due to high daily variability in connectivity data. Connectivity matrices used in this investigation have large relative standard deviations for larval transition probabilities between reefs over the 24 days used (SFigure 8). Using alternative summarisation methods that capture ecological-timescale patterns in coral larval dispersal in future investigations, such as cumulated implicit connectivity (Vogt-Vincent et al., 2023), may find increased importance of both depth and reef connectivity for resilience, in line with other studies (Hock & Mumby, 2017; Torres et al., 2024). Choukroun et al (2025) performed a comparison of three GBR-scale connectivity models, including GBR1, and suggest the use of a model ensemble approach to estimate larval dispersal due to the high sensitivity of connectivity predictions to model specification. Further improvements and research in connectivity modelling are necessary when incorporating connectivity as a supplementary factor in reef resilience and intervention-planning analyses.

Under all GCMs many reefs are projected to suffer substantial declines in coral cover relative to 2025 levels, with an average of 38% of reefs (924 reefs) clustered into the Low cover cluster at the management area scale across GCMs. NorESM2-MM and ACCESS-ESM1-5 GCMs convey the most coral cover maintenance into the future. Differences between Low, Medium and High reef clusters in cover time series are most notable under these two GCMs (Figure 3; STable 2). Maintenance of coral cover into the future is linked to the Equilibrium Climate Sensitivity of the GCM used as levels of future recovery and timespan of cover maintenance increase as GCM ECS increases (Figure 3). Under high-ECS GCMs (GFDL-CM4, EC-Earth3-Veg and ACCESS-CM2) large declines and lower rates of recovery in high cover clusters are common compared to lower-ECS GCMs.

Large differences in cover time series seen across the GCMs suggest it is important to consider uncertainty in future conditions when assessing possible biological responses to change. This is supported by recent literature suggesting an effective approach to capturing climate change uncertainty is to model responses using GCMs separately (Shepherd et al., 2018). Despite large differences in coral cover across GCMs, the patterns identified between low, medium and high clusters of reefs suggest that the relative importance of environmental and biological factors such as depth refugia, connectivity and surface DHW levels is consistent across scenarios.

Reef managers may consider the potential for positive flow-on effects due to outgoing connectivity when selecting intervention sites. The necessary connectivity data, however, must first be available to provide a sufficient degree of confidence to support such site selection decisions (compare the ongoing discussion in Mumby et al. (2018), Bode et al. (2018) and Choukroun et al. (2025)). Given connectivity is a transient, and much more difficult, property to measure that is subject to comparatively higher sources of uncertainty, depth may be a more reliable indicator of long-term resilience, particularly where no further information is available.

### Limitations

There are multiple caveats in how coral ecosystem dynamics are modelled in this investigation. Coral colonies are modelled as diameter values that increase over time through growth. Additionally, the influence of light intensity on coral growth or the negative influence of depth on coral growth rates due to reduced available light levels are not modelled in this investigation (Baker & Weber, 1975; Huston, 1985; Ruiz-Diaz et al., 2022). If included in this investigation, it is possible that reef recovery ability, via regrowth, would be reduced at greater depths, reducing the positive effect of depth refugia for temporal resilience. Additionally, reduced growth rates at increased depths may reduce carbonate budget maintenance due to lower rates of CaCO_3_ accretion. Due to the availability of quality future projected data this investigation does not consider other important impacts such as waves, cyclones, ocean currents and Crown of Thorns Starfish (CoTS) on coral colony survival and bleaching (Emslie, Logan, et al., 2024; Emslie, Ceccarelli, et al., 2024). Uncertainty in connectivity modelling outputs, satellite-derived bathymetry data and future thermal stress projections likely influence the results obtained in this investigation. While the resolution of connectivity data used in this investigation improves upon data used in comparable studies (Bozec et al., 2025), the time-mean implementation used may not effectively capture long term patterns (Vogt-Vincent et al., 2023). Using alternative connectivity implementation methods that better capture the ecological-timescale trends in reef connections rather than the high daily variability, would improve confidence in the connectivity findings of this investigation (Vogt-Vincent et al., 2023). Increased samples to inform baseline connectivity data, along with the derivation of connectivity from an ensemble of biophysical models, would also improve confidence in connectivity data for ecological modelling and management applications (Choukroun et al., 2025).

DHW data used in this investigation have not been bias corrected for the Great Barrier Reef region and are part of a wider Australian dataset (Dixon et al., 2022) developed using statistical downscaling. Lachs et al. (2021) suggest the utility of DHW levels for prediction of coral bleaching can be improved by reducing the threshold for HotSpot Anomaly values from MMM +1 to MMM and reducing the accumulation window from 12 weeks to 8 weeks. The presented analyses assessed a range of future climate responses, thermal projections and ecological parameterisations of ADRIA-CoralBlox to account for many possible futures and suggests the factors that influence reef resilience are consistent across these scenarios.

Highly accurate and precise estimates of reef depth across the GBR are currently not possible to obtain with (publicly) available data. Data adopted for the given assessment are said to be accurate up to 15m from Mean Sea Level (GBRMPA, 2020; Li et al., 2021), however there are many sources of uncertainty in the depth estimates used for this study. Satellite-derived estimates are subject to different bottom types (rock, sand, seagrass, etc) and their effect on reflectance, the water penetration depth of satellite sensors, influence of local conditions (e.g., turbidity and wave activity), among many other factors. The median depths for each indicative reef area were extracted from bathymetry data using zonal statistics and depth values beyond the stated 15m accuracy threshold (GBRMPA, 2020) were included. The resulting depth estimates are therefore highly uncertain and are only adopted as a useful indication for the purpose of the presented analysis. They are not intended to be accurate and precise reef depths across the GBR.

In addition to the usual challenges of deriving data from satellite imagery, it is acknowledged that there is high potential for misalignment between the bathymetry data and GBRMPA reef outlines. Hammerton and Lawrey (2024), for example, identified 120 false positives, where features identified as reefs were sand banks or rocky reefs (rather than coral reefs). Reefs with bathymetry data covering less than 5% of their indicative area were excluded from analyses (see SFigure 2). Existing GBRMPA reef outlines were adopted to align with other ongoing modelling and analyses conducted as part of the Reef Restoration and Adaptation Program (https://gbrrestoration.org, RRAP, 2025).

The modelling conducted with ADRIA-CoralBlox itself is subject to many uncertainties. For one, a formal mechanistic and spatial validation of the model represents an area for future refinement and is the focus of a future publication. The consistency of projected patterns with known coral-climate relationships and reef characteristics provides confidence in the model’s ability to meaningfully capture ecological signals at the GBR-scale. The mechanistic basis of the model supports the robustness of these findings for understanding broad-scale coral cover responses to climatic drivers. The aim here, however, was to assess the relative influence of reef characteristics on reef resilience across a range of future conditions and parameterisations, rather than provide spatially explicit and precise predictions of reef outcomes. Using this approach, results suggest reef depth is an important factor for reef resilience under thermal stress.

Improvements in remote sensing approaches for broad scale bathymetry measurement and understanding of how depth impacts experienced thermal stress on reefs will further enhance investigations of depth as a critical factor for reef resilience. Additional analyses could build on the findings of this study by assessing the impacts of prioritising reef depth for reef intervention decision outcomes. Future analyses could incorporate other environmental and ecological stressors such as wave activity, ocean acidification, and Crown-of-Thorn Starfish (Eyre et al., 2018; Grimaldi et al., 2022, 2023; Lin et al., 2023; D. Liu et al., 2024; McClanahan et al., 2005).

## Conclusion

The results presented suggest that when modelled as part of a large and complex ecosystem, reef depth is a key factor contributing to ongoing reef resilience. This investigation provides evidence in support of depth as a form of thermal refugia for shallow-water reefs on the GBR. Targeting reefs with median depths greater than 10m may aid in identifying the most resilient reefs under climate change. Further analyses that investigate the benefits of incorporating depth-based resilience into reef intervention decision making, as well as further research and development in connectivity modelling methods for whole-ecosystem scales are recommended.

## Acknowledgements

The authors acknowledge the Traditional Owners of the Sea Country on which the modelling was conducted for. The authors additionally acknowledge the Bindal, Jagera, and Ngunnawal Peoples, the lands on which the authors conducted the model development, analysis, and writing. The authors would like to acknowledge and thank Shaun Wilson for comments and discussion during manuscript revision. The authors also acknowledge the contributions of Barbara Robson and Jessica Benthuysen for comments provided early in the manuscript preparation. The authors would like to additionally acknowledge and thank Shaun Wilson and Juan Carlos Ortiz for discussions around carbonate budget thresholds.

This research was funded by the Reef Restoration and Adaptation Program and facilitated by the Australian Institute of Marine Science. The authors acknowledge thermal projection data sourced from HighResCoralStress datasets, larval transition probability data derived from the eReefs GBR1 hydrodynamic model, and spatial and depth data available from the Great Barrier Reef Marine Park Authority. The eReefs Project is a public-private collaboration between Australia’s leading operational and scientific research agencies, government, and corporate Australia.

## CRediT Statement

**Benjamin Grier:** conceptualisation, data curation, formal analysis, investigation, methodology, software, visualization, writing – original draft, writing – review and editing. **Takuya Iwanaga:** conceptualisation, data curation, formal analysis, investigation, methodology, software, visualization, writing – original draft, writing – review and editing. **Pedro Ribeiro de Almeida:** data curation, software, formal analysis, writing - methodology, writing - review and editing, visualisation. **Chinenye Ani:** data curation, writing - methodology, writing - review and editing. **Sam Matthews:** Writing - review and editing.

## Data Availability Statement

The full reproducible investigation, and links to required data sources, can be found at Grier and Iwanaga (2025, https://doi.org/10.5281/zenodo.17333564).

## Ethics Statement

The presented work did not involve animal or human participants, or personal and confidential data.

