## Supplemental Material for "Importance of depth refugia for reef resilience and intervention on the Great Barrier Reef under future climate change"

### Supplementary Material

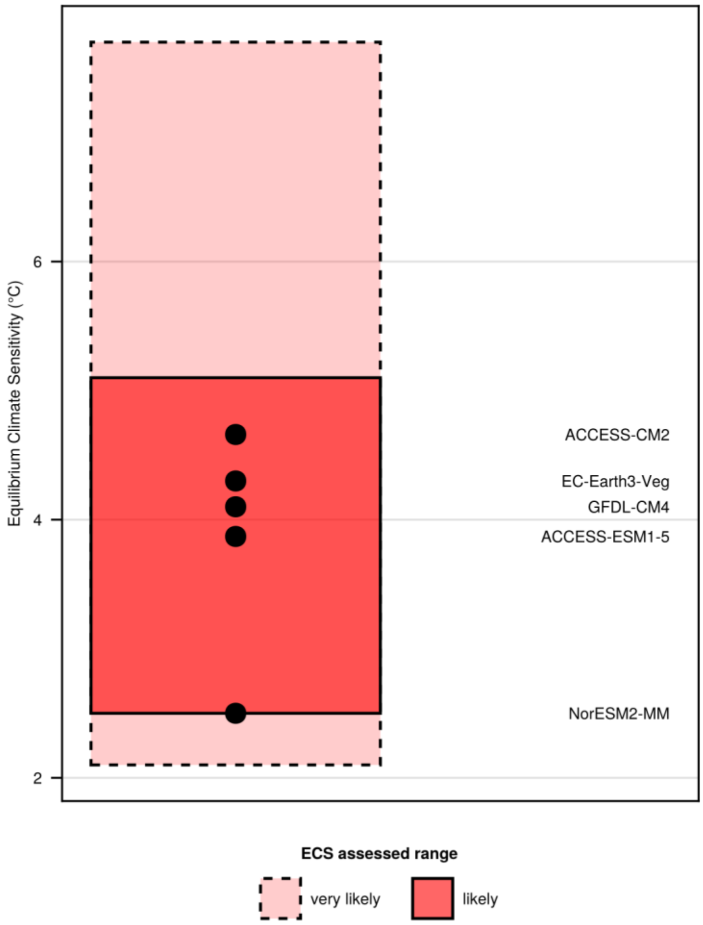


*SFigure 1: Equilibrium Climate Sensitivity (ECS) values in °C for the GCMs selected (point values). The pink - dashed box represents the high-certainty 'very likely' range for ECS values and the solid red box represents low-certainty 'likely' range for ECS values (Forster et al., 2021).*

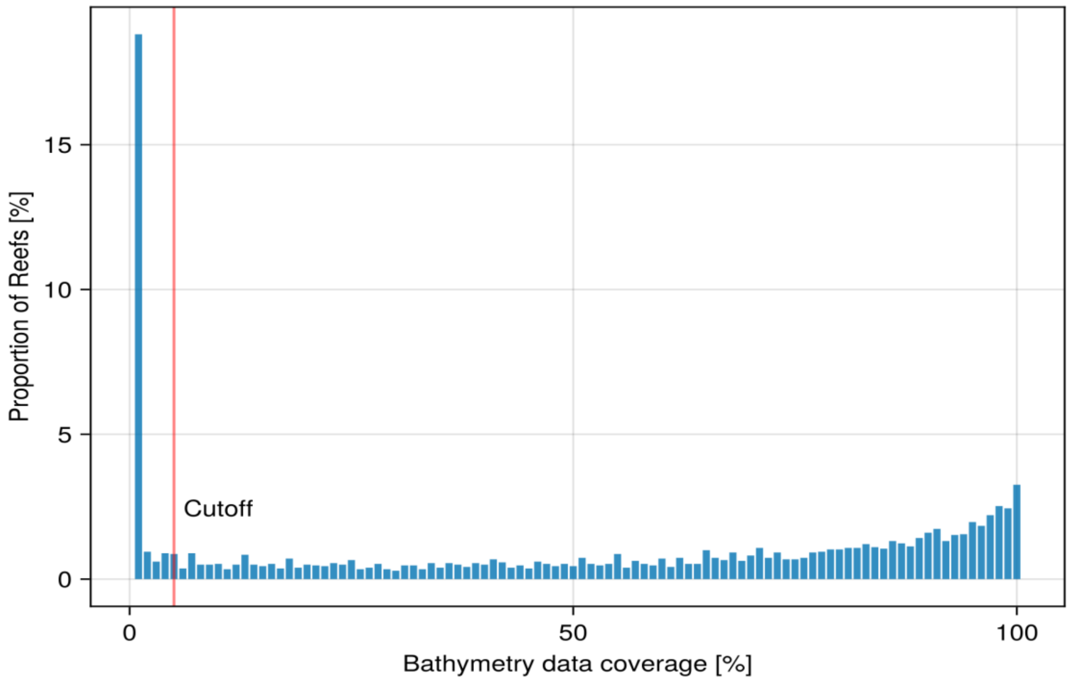


*SFigure 2: Distribution of GBRMPA (2020) satellite derived depth data coverage levels across 3806 possible reefs. Reefs below the 5% data coverage threshold (represented by the red vertical line) were removed from the investigation.*

3

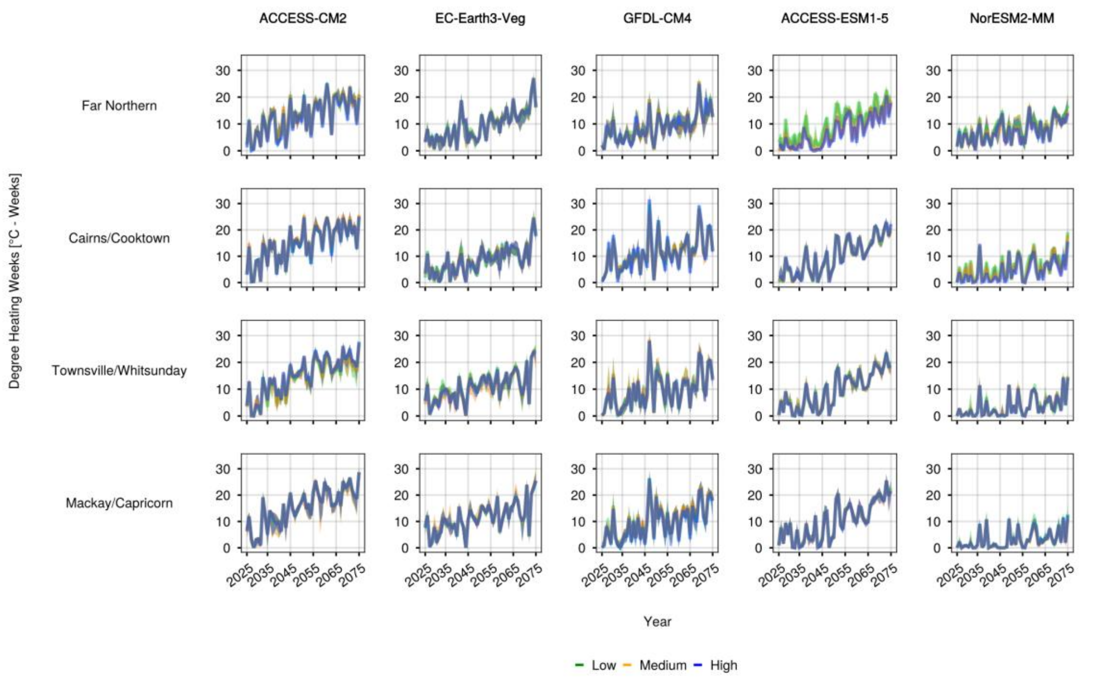


*SFigure 3: Reef Degree Heating Week time series clustered within each management area, for each GCM analysed. Time series were clustered based on their behaviour between 2030 and 2060. Time series with comparatively low median area-relative*

*cover is indicated in green, intermediate cover in orange, and high in blue. Solid lines represent the cluster median time series and bands represent the 2.5 to 97.5 percentile confidence intervals. Each management area is represented five times (for each GCM), with each GCM displayed horizontally.*

4

| Metric | Used in | Definition | Justification |
|---|---|---|---|
| Percentage coral cover time series | Time series clustering, Carbonate budget analysis | Total coral cover as a percentage of reef area | To summarise coral population health over time. |
| Median depth | Time series cluster assessment Carbonate budget analysis | Reef median depth as calculated with zonal raster analysis | Depth is a critical form of environmental refugia from high thermal stress |
| Mean time series DHW | Time series cluster assessment | Reef mean DHW value across time series | To quantify the surface thermal stress experienced on each reef over time series. |
| Weighted incoming connectivity | Time series cluster assessment, Carbonate budget analysis | Incoming connectivity strength weighted by mean time series coral cover of source reefs | To approximate the larval flow received by a reef over the projected time series. |
| Outgoing connectivity strength | Time series cluster assessment, Carbonate budget analysis | Sum of a reef's outgoing larval transition probability values. | To understand the contribution of a reef to the connectivity network and the possibility for flow-on effects from reef states. |
| Larval retention probability | Time series cluster assessment | Diagonal larval transition matrix values. | To understand the relative proportion of larvae retained on a reef. |
| Carrying capacity | Time series cluster assessment | Reef habitable area [$m^2$] | Captures the size of the viable reef extent. |

*STable 1: Quantitative metrics used in this investigation.*

| GCM | Management area | CID - cover | CID - DHW |
|---|---|---|---|
| ACCESS-CM2 | Cairns/Cooktown | 24.42 | 1.99 |
| ACCESS-ESM1-5 | Cairns/Cooktown | 58.13 | 6.0 |
| EC-Earth3-Veg | Cairns/Cooktown | 57.65 | 8.05 |
| GFDL-CM4 | Cairns/Cooktown | 14.82 | 10.44 |
| NorESM2-MM | Cairns/Cooktown | 124.38 | 20.99 |
| ACCESS-CM2 | Far Northern | 45.04 | 8.11 |
| ACCESS-ESM1-5 | Far Northern | 114.53 | 35.96 |
| EC-Earth3-Veg | Far Northern | 89.53 | 4.2 |
| GFDL-CM4 | Far Northern | 50.46 | 10.25 |
| NorESM2-MM | Far Northern | 64.85 | 11.11 |

| | | | |
|---|---|---|---|
| ACCESS-CM2 | Mackay/Capricorn | 63.22 | 2.27 |
| ACCESS-ESM1-5 | Mackay/Capricorn | 80.88 | 2.5 |
| EC-Earth3-Veg | Mackay/Capricorn | 64.66 | 1.81 |
| GFDL-CM4 | Mackay/Capricorn | 44.84 | 4.67 |
| NorESM2-MM | Mackay/Capricorn | 96.25 | 1.73 |
| ACCESS-CM2 | Townsville/Whitsunday | 63.19 | 7.67 |
| ACCESS-ESM1-5 | Townsville/Whitsunday | 67.85 | 3.24 |
| EC-Earth3-Veg | Townsville/Whitsunday | 64.37 | 5.32 |
| GFDL-CM4 | Townsville/Whitsunday | 50.4 | 3.43 |
| NorESM2-MM | Townsville/Whitsunday | 101.68 | 2.98 |

*STable 2: Complexity invariant distance (CID) between low and high cluster median time series in each management area under each GCM. CID values are calculated for percentage cover time series (Figure 3) and Degree Heating Week (DHW) time series (SFigure 3). Complexity invariant distance represents the Euclidean distance between time series, corrected by the complexity of the two time series. CID values are used to quantify overall differences between these two cluster levels at management area and GCM levels. CID values for percentage cover and DHW time series are not directly comparable as the time series use different units.*

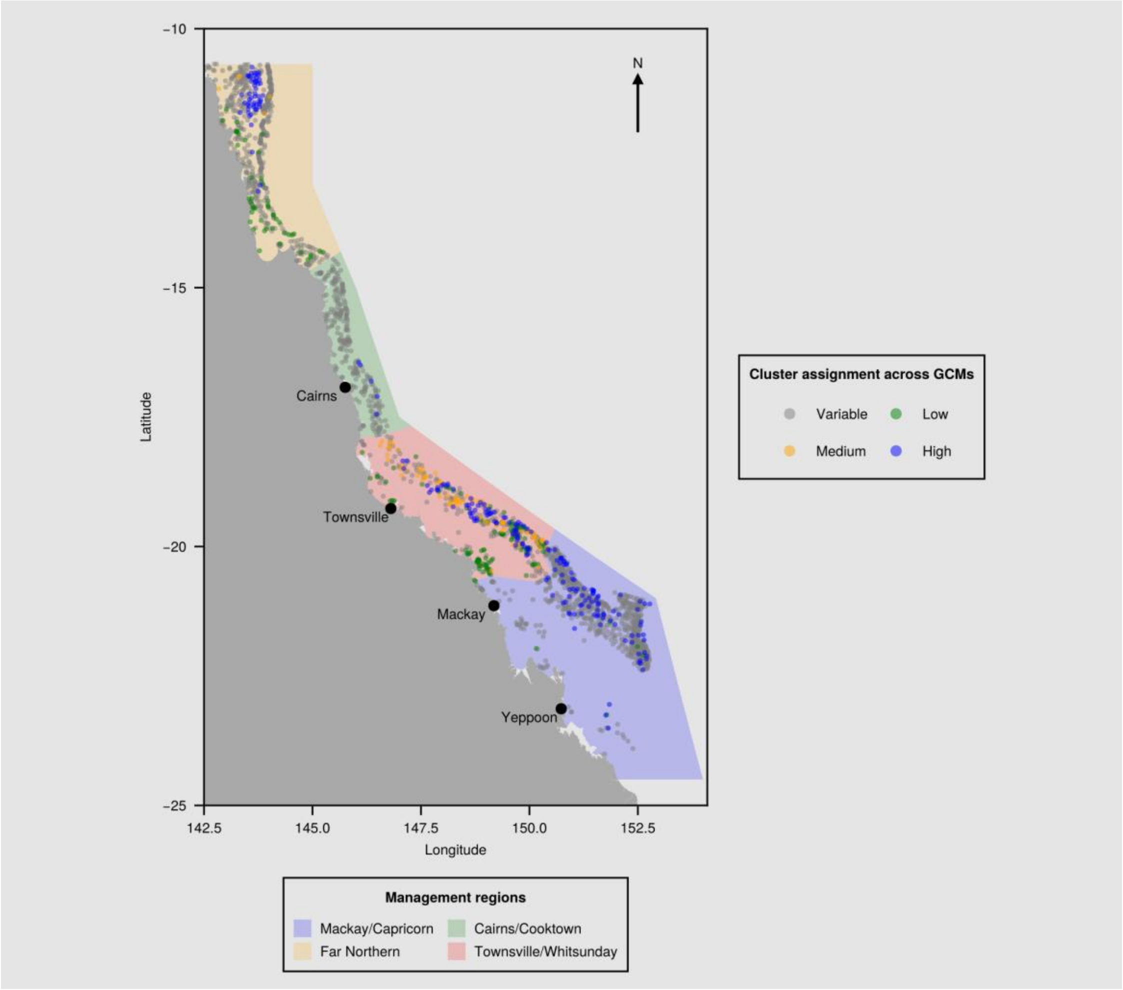

*SFigure 4: Map of the Great Barrier Reef region, including GBRMPA management areas (polygons) and all considered reefs (points), coloured by management area scale cluster assignment. Colour-coded areas outline GBR marine park management regions, where orange represents the Far Northern management region, green represents Cairns/Cooktown, red covers Townsville/Whitsunday and blue covers Mackay/Capricorn (GBRMPA, 2007). Reef centroids are marked individually for visual clarity. Reefs that do not have a consistent cluster assignment level across all GCMs at management area scale are coloured grey. All consistently clustered reefs are coloured according to their management area scale cluster assignment. Green represents reefs in the low cover cluster, orange represents reefs in the medium cover cluster and blue represents reefs in the high cover cluster. All 2197 reefs included in this investigation are represented in this figure. Annotations represent major coastal settlements along the GBR.*

5

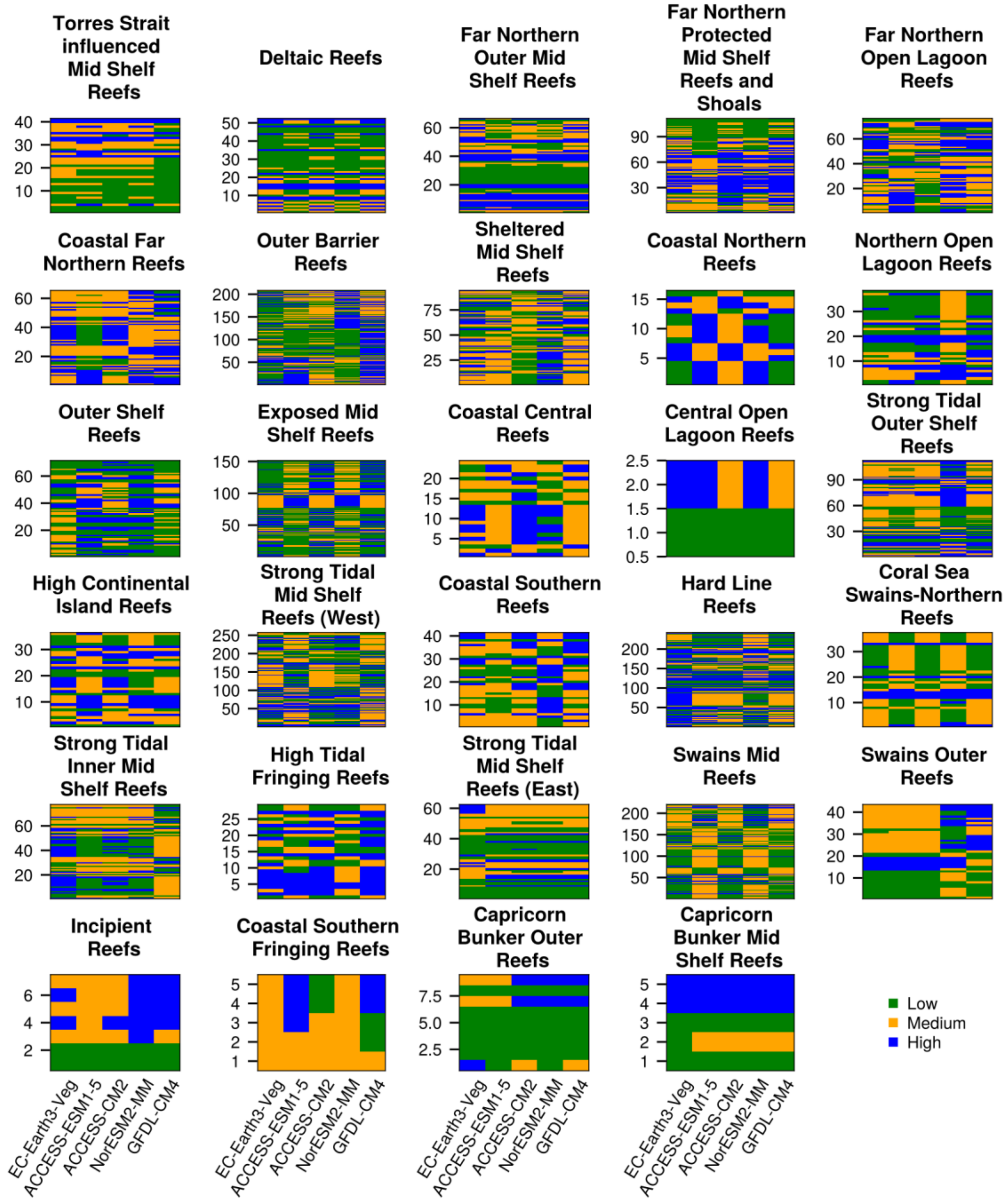


*SFigure 5: Heatmaps indicating reef cluster assignments within each bioregion across the five GCMs. Green indicates a low cover cluster assignment, orange represents a medium cover assignment and blue represents assignment of a reef to a high coral cover cluster. The vertical width of reef elements varies depending on the number of reefs within the bioregion (displayed on the y axis). The x axis represents the different GCM levels (EC-Earth3-Veg, ACCESS-ESM1-5, ACCESS-CM2, NorESM2-MM and GFDL-CM4).*

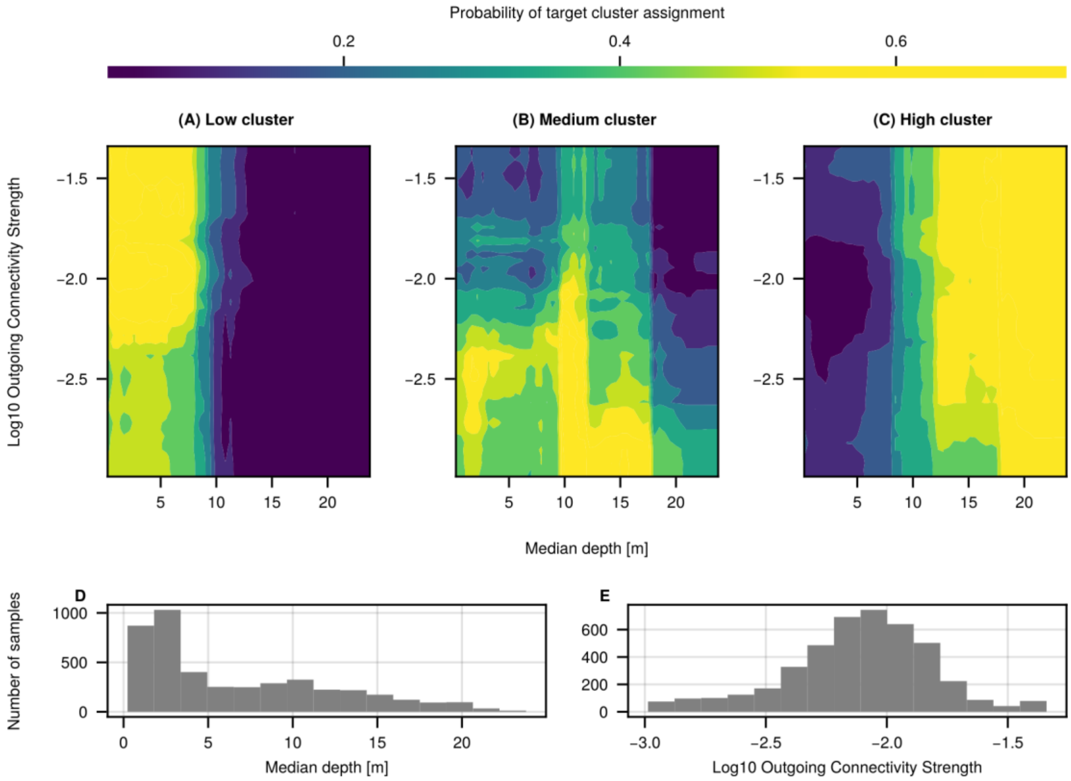


*SFigure 6: Partial dependence plot indicating the probability of cluster assignment along interacting values of median reef depth (meters) and log10 weighted outgoing connectivity. Colour indicates the probability of assignment to each cluster, with lighter colours indicating higher probability of assignment. Panel A shows the probability of assignment for the Low cover cluster, Panel B shows Medium cover cluster and Panel C shows assignment probability for the High cover cluster. Probability values for each cluster are generated by the random forest classifier using each combination of depth and weighted outgoing connectivity values along the axes at 50 grid points. The actual distributions of depth and outgoing connectivity values for reefs are displayed in panels D and E respectively.*

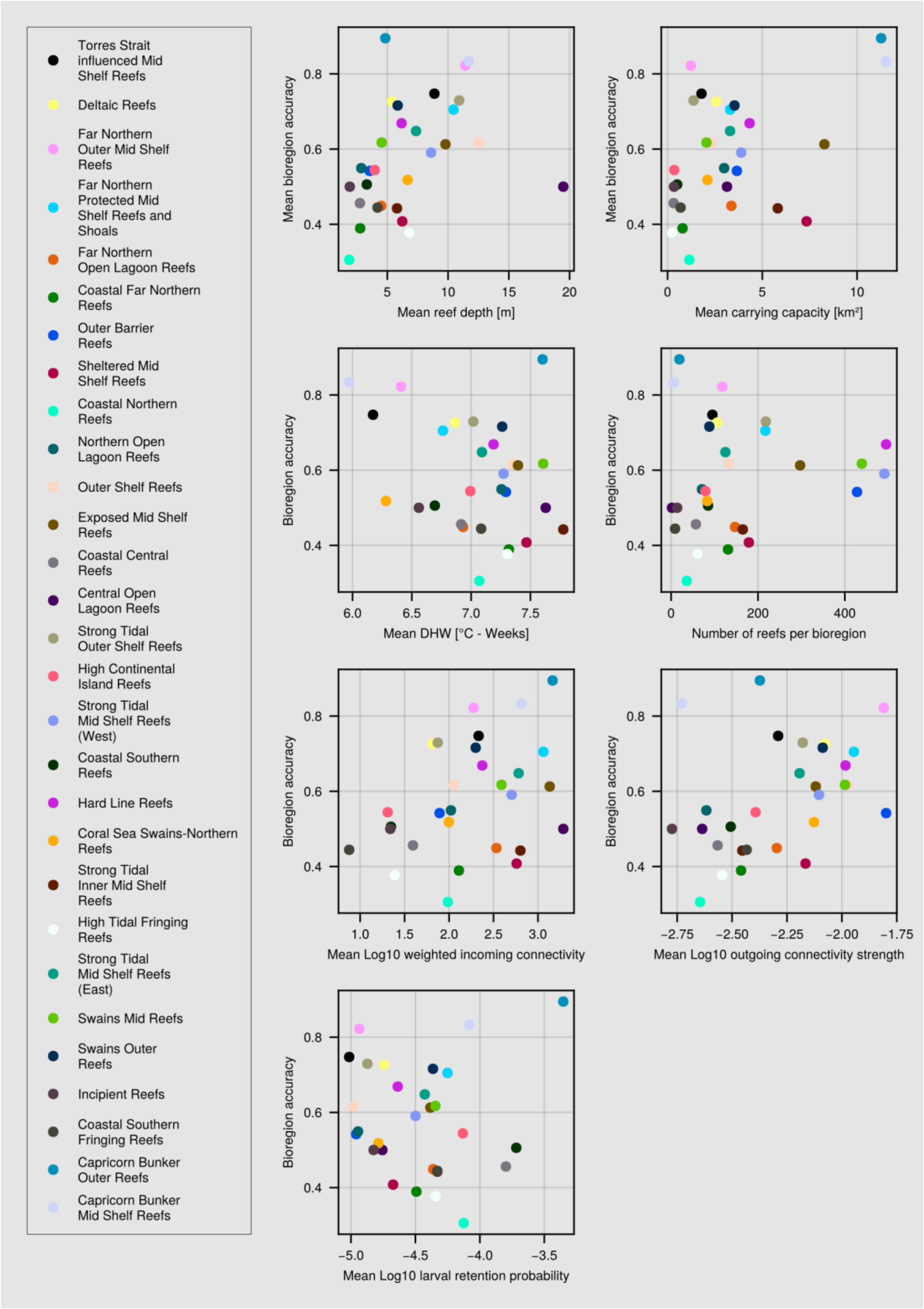


*SFigure 7: Scatter plots indicating the bioregion test-prediction accuracy (proportion of reefs in each bioregion assigned correct clusters) from the Random Forest model. Plot panels represent the bioregion accuracy across various predictor ranges and bioregion attributes, such as mean bioregion depth, carrying capacity, DHW value, total connectivity strength and the number of reefs in each bioregion.*

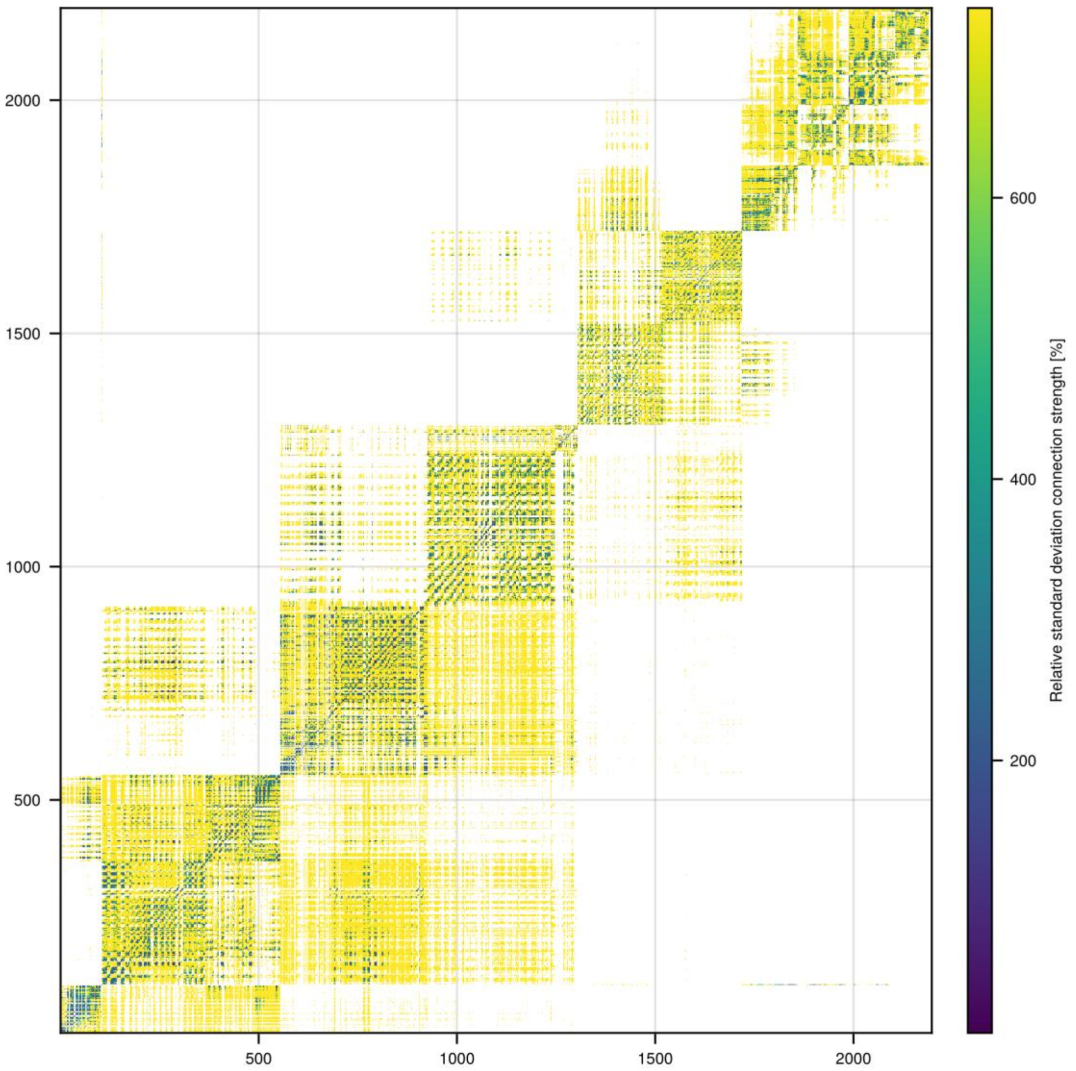


*SFigure 8: Heatmap indicating reef connectivity relative standard deviation (RSD %). Reef connectivity relative standard deviation represents the reef standard deviation connection strength as a percentage of reef mean connection strength. Lighter colours indicate higher RSD % values.*

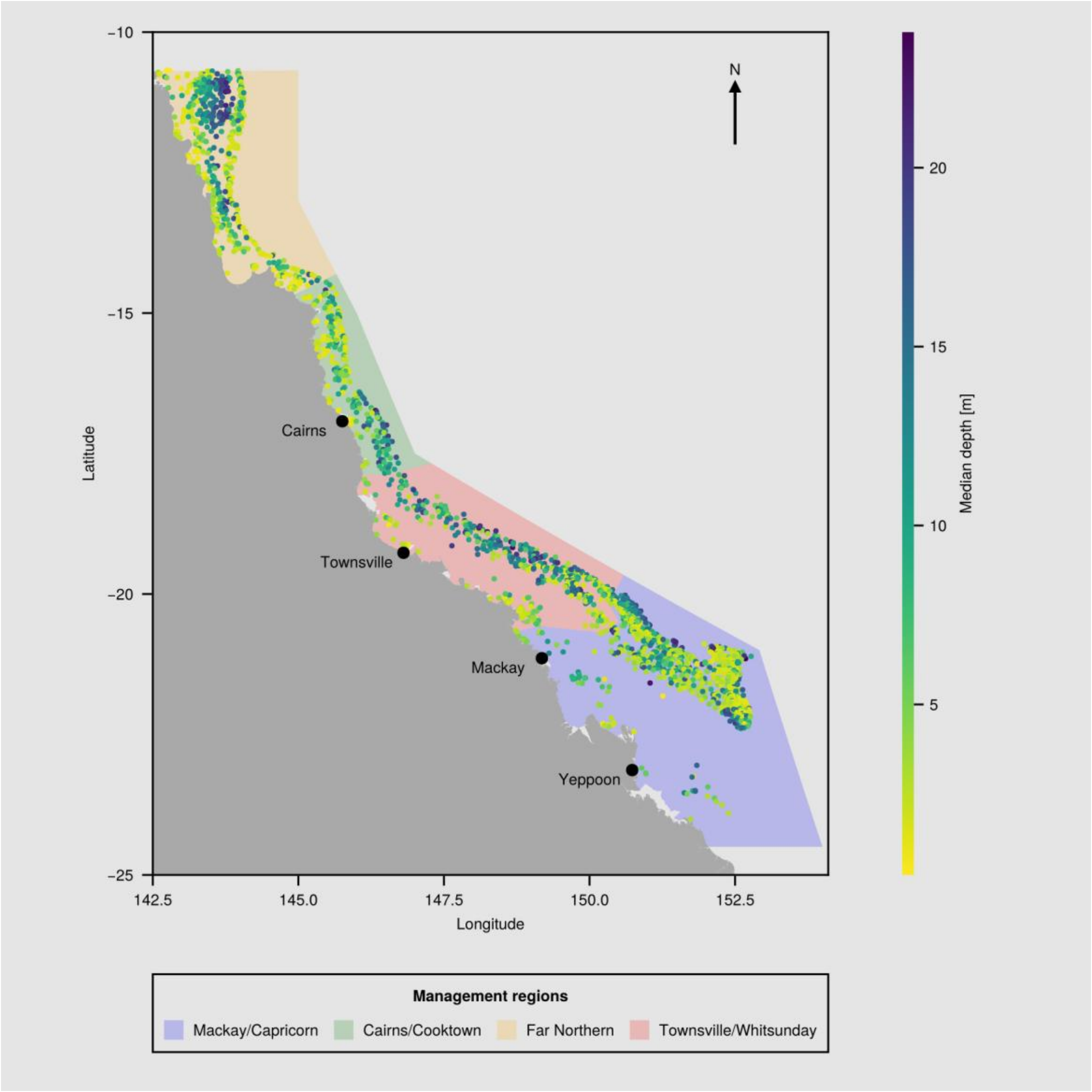


*SFigure 9: Map of the Great Barrier Reef region, including GBRMPA management areas (polygons) and all considered reefs (points), coloured by median reef depth. Colour-coded areas outline GBR marine park management regions, where orange represents the Far Northern management region, green represents Cairns/Cooktown, red covers Townsville/Whitsunday and blue covers Mackay/Capricorn (GBRMPA, 2007). Reef centroids are marked individually for visual clarity. Darker colours represent greater reef depth values than lighter colours. All 2197 reefs included in this investigation are represented in this figure. Annotations represent major coastal settlements along the GBR.*

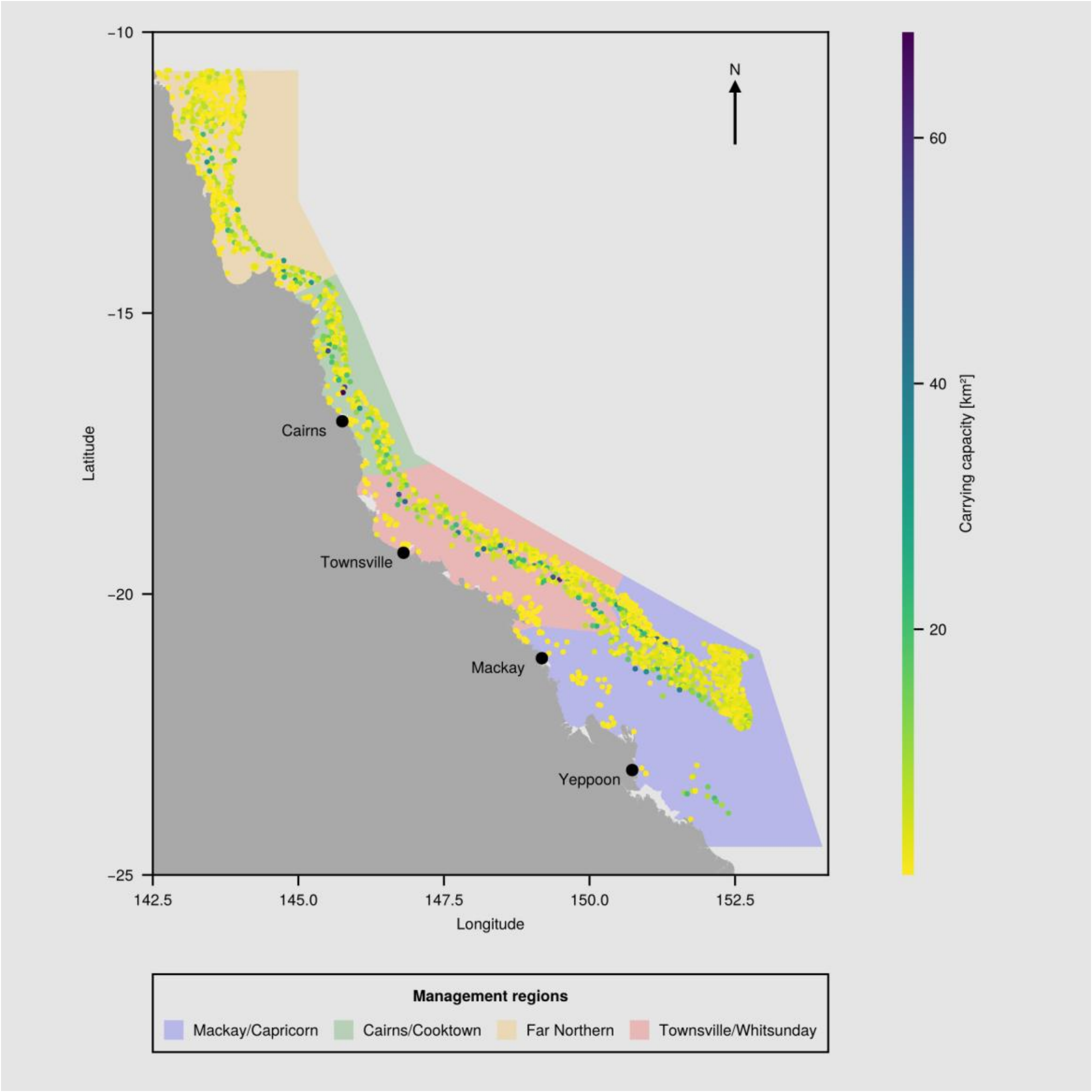


*SFigure 10: Map of the Great Barrier Reef region, including GBRMPA management areas (polygons) and all considered reefs (points), coloured by reef carrying capacity (coral habitable area). Colour-coded areas outline GBR marine park management regions, where orange represents the Far Northern management region, green represents Cairns/Cooktown, red covers Townsville/Whitsunday and blue covers Mackay/Capricorn (GBRMPA, 2007). Reef centroids are marked individually for visual clarity. Darker colours represent greater carrying capacity area values than lighter colours. All 2197 reefs included in this investigation are represented in this figure. Annotations represent major coastal settlements along the GBR.*

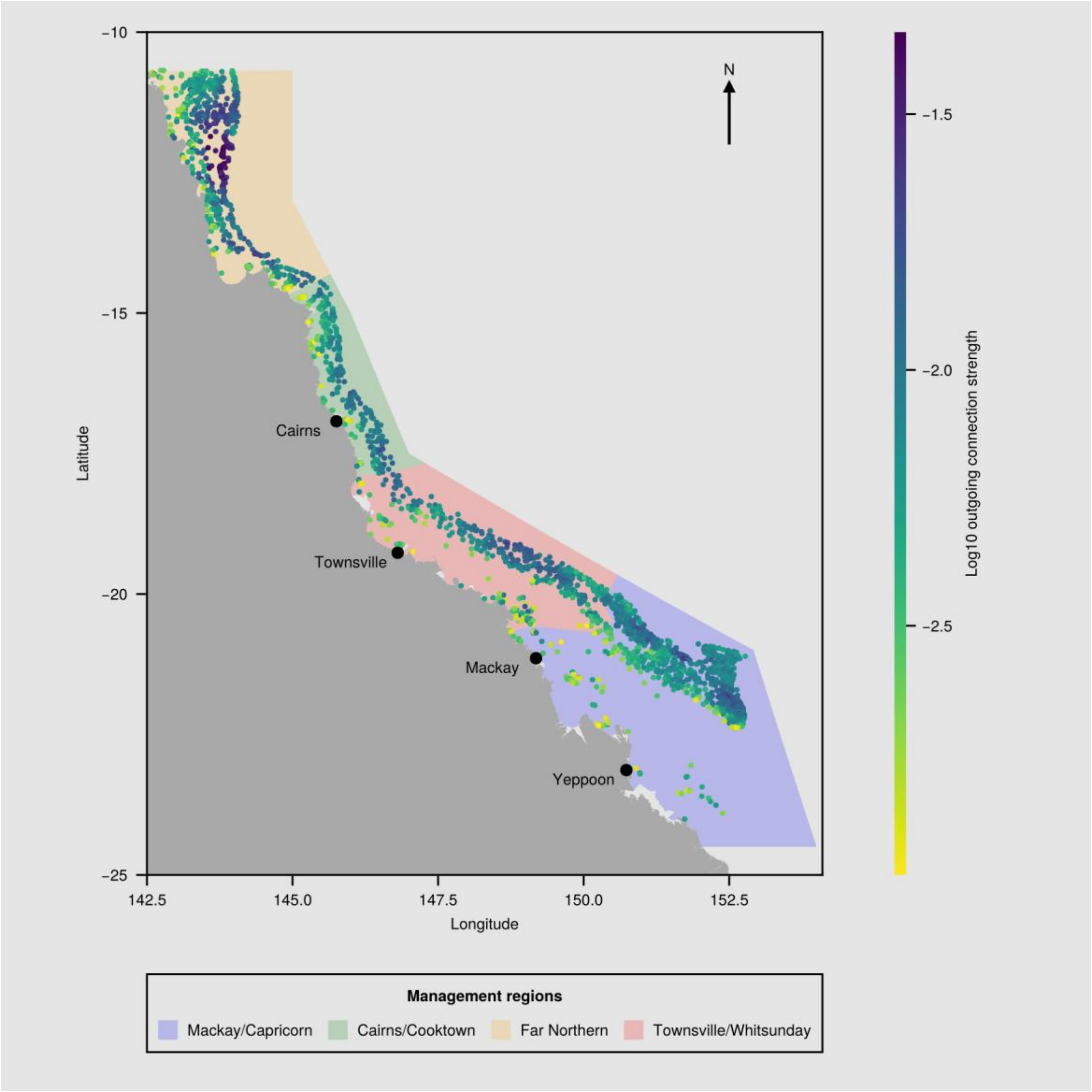


*SFigure 11: Map of the Great Barrier Reef region, including GBRMPA management areas (polygons) and all considered reefs (points), coloured by outgoing connection strength (log10). Colour-coded areas outline GBR marine park management regions, where orange represents the Far Northern management region, green represents Cairns/Cooktown, red covers Townsville/Whitsunday and blue covers Mackay/Capricorn (GBRMPA, 2007). Reef centroids are marked individually for visual clarity. Darker colours represent higher outgoing connection strength values than lighter colours. All 2197 reefs included in this investigation are represented in this figure. Annotations represent major coastal settlements along the GBR.*

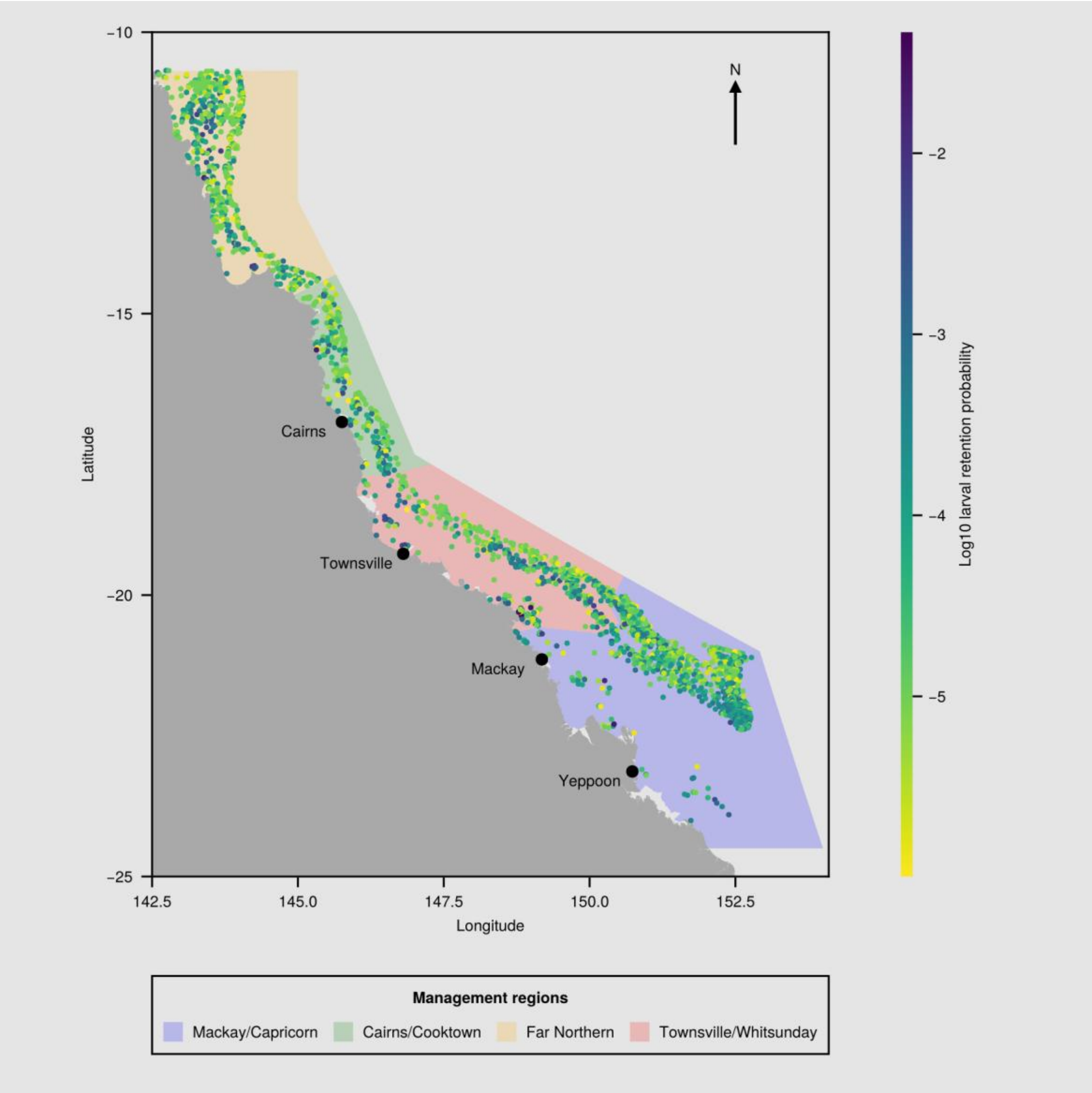


*SFigure 12: Map of the Great Barrier Reef region, including GBRMPA management areas (polygons) and all considered reefs (points), coloured by larval retention probability (log10). Colour-coded areas outline GBR marine park management regions, where orange represents the Far Northern management region, green represents Cairns/Cooktown, red covers Townsville/Whitsunday and blue covers Mackay/Capricorn (GBRMPA, 2007). Reef centroids are marked individually for visual clarity. Darker colours represent higher probability of retaining spawned larvae than lighter colours. All 2197 reefs included in this investigation are represented in this figure. Annotations represent major coastal settlements along the GBR.*

#### Degree Heating Week data processing

Monthly mean SST climatological baselines, for each 1km$^2$ pixel, were calculated from the daily SST data following NOAA methodology (G. Liu et al., 2017; Skirving et al., 2020). For each pixel of SST data, the Monthly Mean SST climatology was calculated by computing the mean SST value for each calendar month (January through December) across the 28-year baseline period from 1985 to 2012. For each month separately, this climatological baseline was adjusted to centre at year 1988.2857, which attempts to account for any long-term trends within the baseline years themselves (see Skirving et al. (2020)). The adjustment ensures that the climatology represents conditions at a specific temporal reference point rather than being influenced by gradual warming or cooling trends that may have occurred during the 1985-2012 period. The resulting monthly climatologies serve as a standardized reference against which current SST conditions can be compared to identify thermal anomalies and assess coral bleaching risk, following the methodology established by Skirving et al. (2020).

Degree Heating Weeks (DHWs) were derived using the “HotSpot SST Anomaly” approach (Skirving et al., 2020). The “HotSpot SST Anomaly” represents thermal stress experienced by coral reefs and is calculated as the difference between daily SST values and the maximum of a pixel’s monthly mean climatology across all twelve months (i.e., the maximum monthly mean SST). This approach uses the annual maximum temperature as the baseline because corals are adapted to withstand their local warmest conditions under normal seasonal cycles. When temperatures exceed this seasonal maximum, corals experience thermal stress that can lead to bleaching. Only positive anomalies, greater than 1°C, are considered relevant for coral stress, as temperatures below the seasonal maximum do not contribute to bleaching risk.

#### ADRIA-CoralBlox

The CoralBlox model incorporates linear extension (growth) and background mortality rates that are dependent on the size and functional group of a coral colony.

Environmental disturbances and reproduction are not directly considered within CoralBlox, and are instead provided by the ADRIA (Adaptive Dynamic Reef Intervention Algorithms) platform (T. Iwanaga et al., 2023). Within ADRIA, coral life history and environmental events are implemented in the following order: coral colony growth; coral reproduction; larval connectivity and settlement; and coral bleaching due to heat stress.

Coral fecundity rates are dependent on colony size and functional group following the results of Hall & Hughes (1996). Coral fertilization rates are not considered within this model, however larval survival and viability are incorporated in connectivity modelling (see Connectivity modelling). Viable larvae settle at a reef location and are incorporated into the model as the smallest size class but are not directly impacted by modelled bleaching mortality. There has been contrasting evidence for juvenile corals being immune to thermal bleaching events (Depczynski et al., 2013), it is assumed the high background mortality of juveniles includes thermal stress (Álvarez-Noriega et al., 2018).

#### Model implementation

ADRIA-CoralBlox implements five functional groups common to the GBR, tabular Acropora, corymbose Acropora, corymbose non-Acropora, small massives and large massives. CoralBlox uses user-defined size classes that influence size-dependent growth and mortality rates, through which the coral colonies progress as they grow in diameter. Size classes were initially informed by Bozec et al. (2022).

The area of each coral colony is tracked in CoralBlox by the area of a circumference. A coral population is then represented by a discretised distribution of $p(x)$ coral diameters, composed of smaller uniform distributions, each representing the number of corals within a certain diameter range:

$$\rho(x) = \begin{cases} \lambda_1, & d_0 < x < d_1 \\ \vdots & \vdots \\ \lambda_N, & d_{N-1} < x < d_N \\ 0, & x \notin [d_0, d_N] \end{cases}$$

where $N$ represents the number of intervals used to discretise $p(x)$, and $d_0$ and $d_N$ are the diameters of the smallest and biggest coral colonies, in that order. The coral cover for the entire population is given by:

$$\begin{aligned} C &= \int_{d_0}^{d_N} \pi\, x^2 \rho(x)\, dx \\ &= \sum_{n=1}^{N} \pi\, \lambda_n \int_{d_{n-1}}^{d_n} x^2\ dx \\ &= \sum_{n=1}^{N} \pi\, \lambda_n \left( \frac{d_n^3 - d_{n-1}^3}{3} \right) \end{aligned}$$

These are uniform distributions that compose $p(x)$, a coral block characteristed by a diameter density $\lambda_n$ (the block's height) and a diameter range $[d_{n-1}, d_n]$ (the base of the block). Therefore, instead of working with a continuous diameter's distribution, CoralBlox updates each coral block separately. A mortality event is modelled as a decrease in a coral block's density, and a growth event as an increase in a coral block's diameter range limits. Both growth and mortality rates are size class dependent, which means the same growth and mortality rates are used for blocks within a size class. SFigure 13 shows a model's behaviour under low disturbance and no interventions.

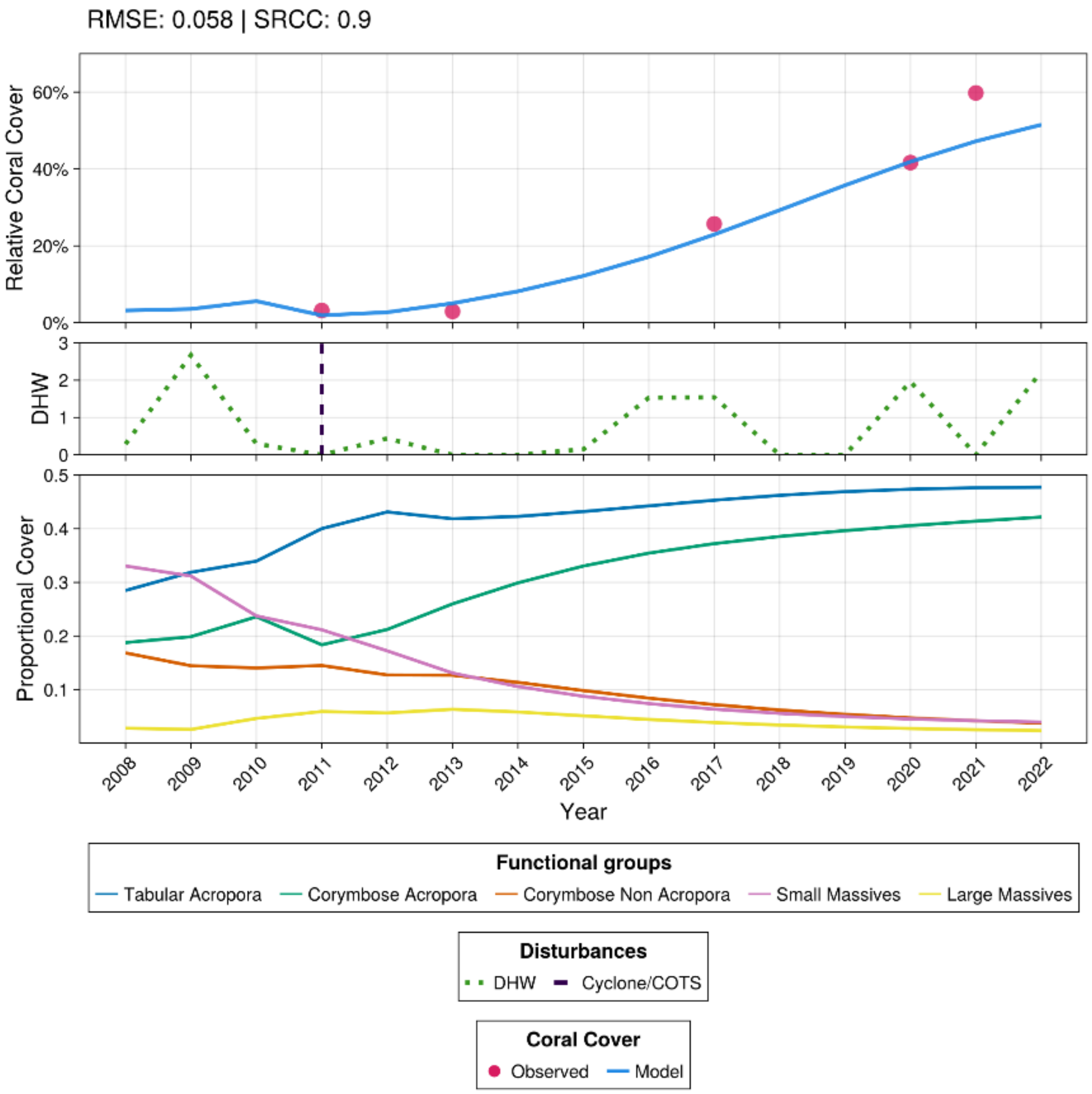


*SFigure 13: Time series comparing modelled and observed relative coral cover for Ben reef showing a recovery process under very low disturbance and no interventions. The observed data used here is from the AIMS Long-Term Monitoring Program (LTMP) . Error metrics including Root Mean Square Error (0.06) and the Spearman Rank Correlation Coefficient (0.9) is also shown. The second plot shows the DHW and Cyclone or COTS events. The third plot shows the benthic composition for each functional group as a proportion of the reef's cover at each year.*

In ADRIA we track the thermal tolerance of each functional group and size class as a truncated normal distribution that is updated in each timestep following growth and recruitment. This distribution determines the proportion of the population on a reef that is susceptible to bleaching (i.e. has a thermal threshold below the experienced DHW value). Starting distributions for thermal tolerance levels are informed by Bairos-Novak et al. (2021) and expert opinion, and are additionally varied in the scenario exploration.

#### Scenario generation

Scenarios were sampled from the range of environmental and ecological factors using the low-discrepancy Sobol' sampling method (Sobol', 1967, 1993) with Owen scrambling (Owen, 1995). A base of 2 and padding of 32 is used for the Owen scrambling. Ecological factors were sampled using a +/- 10% range relative to their nominal (assumed best guess) value.

#### Sensitivity analysis

A global sensitivity analysis was performed using Shapley Effects (Goda, 2021) implemented in the SAShE.jl package (Ribeiro de Almeida, 2025). For this analysis, 10,000 samples were simulated for a single reef and timestep. Two metrics were considered, difference in coral cover and difference in evenness. The factors analysed were heritability, mean colony diameter, linear extension, mortality base rate, mean and standard deviation of the tolerance distribution, linear extension scale factor, mortality rate scale factor reef habitable area, reef depth, DHW, and initial coral cover. SFigure 14 shows the result of such analysis for different ranges of DHW. In both cases a similar effect takes place: when DHW = 0, initial cover is the most influential factor then, when DHW > 0, depth and DHW start to appear as influential factors and, as DHW values get higher, depth becomes more influential in detriment of DHW.

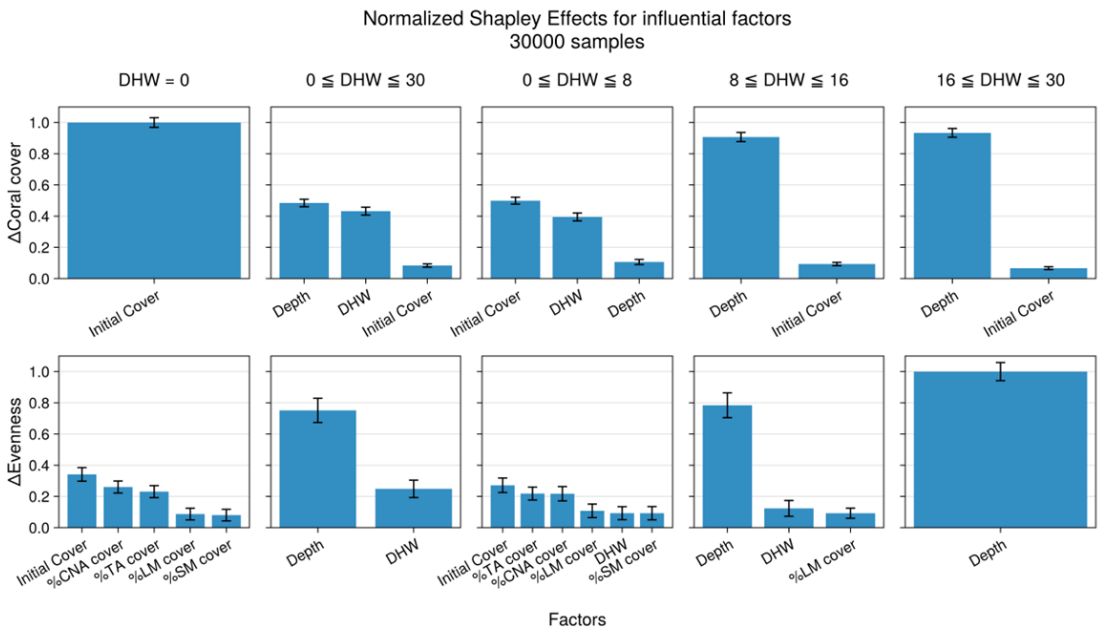


*SFigure 14: Parametric sensitivity analysis conducted with Shapley effects. Each column reflects the sampling of different DHW ranges. The first column represents model runs with DHW = 0, and the remaining columns represent model runs with DHW valued between 0 and 30, 0 and 8, 8 and 16, 16 and 30. The first row shows the Shapley effect for relative changes in coral cover and the second for relative changes in evenness of species abundance. The five functional group names were abbreviated for readability as TA (tabular Acropora), CA (corymbose Acropora), CNA (corymbose non-Acropora), SM (small massives) and LM (large massives).*

#### Time series clustering

The CID time series clustering metric corrects the Euclidean distance between pairs of time series with their relative complexity (which is the quotient of their variability). This is to ensure reefs are clustered based on their underlying temporal dynamics rather than aggregating the coral cover across all timesteps. The process creates a $n * n$ distance matrix (where $n$ is the number of time series in the group) that is used as an input to a k-medoids clustering algorithm (cf. Steinmann et al., 2020). The three reef clusters are labelled with qualitative-ordered labels from Low, Medium and High based on their median cover relative to each other cluster over the 2030-2060 period. The selected time frame prevents the classification from being overly influenced by any steep declines in the initial years of simulations and avoids the post-2060 time period where reefs are projected to be under severe stress and in a declined state (Bozec et al., 2025).

#### ACCESS-ESM1-5 results

ACCESS-ESM1-5 was selected for GCM-level results as an intermediate severity GCM, with an ECS value closest to the midpoint of the IPCC likely range of ECS values (3.87 °C). The following section includes supplemental results for analyses under this GCM. A comparison of results across GCMs can be found in the main text, and individual GCM-level results for other GCMs can be found at Grier and Iwanaga (2025).

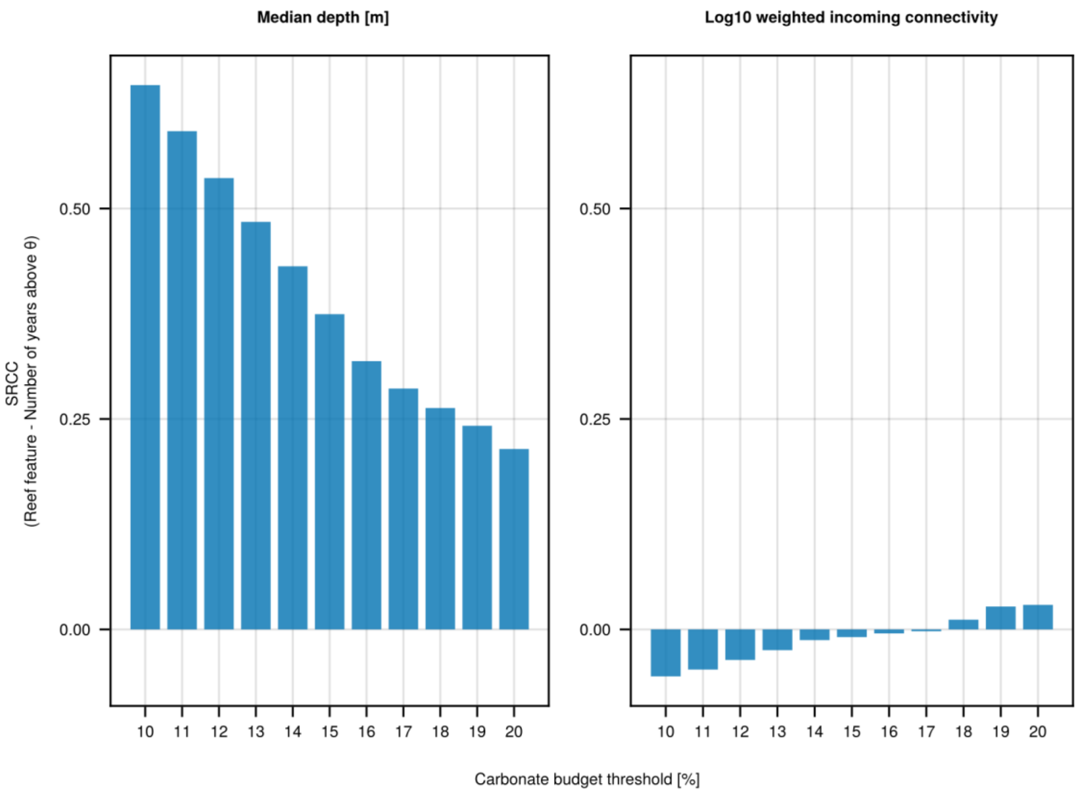


*SFigure 15: Bar plot indicating the Spearman's rank correlation values for the number of years above carbonate budget threshold* $(theta)$ *and, median reef depth and log10 weighted incoming connectivity values. Depth correlation values are displayed in the left plot, while connectivity correlation values are displayed in the right plot. Displays results for ACCESS-ESM1-5 GCM.*

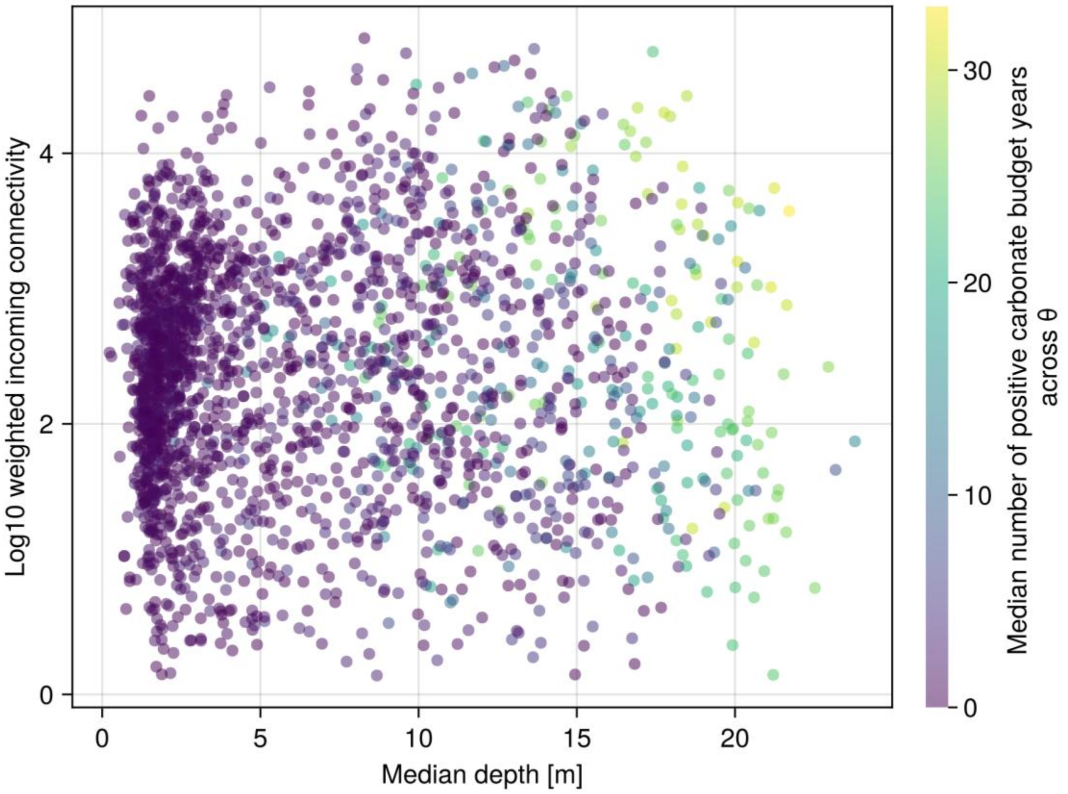


*SFigure 16: Scatter plot indicating reef median depth (metres) and log10 weighted incoming connectivity for each reef. Point colours represent the median number of years each reef maintains positive carbonate budgets across θ values from 10-20% live coral cover. Displays results for ACCESS-ESM1-5 GCM.*

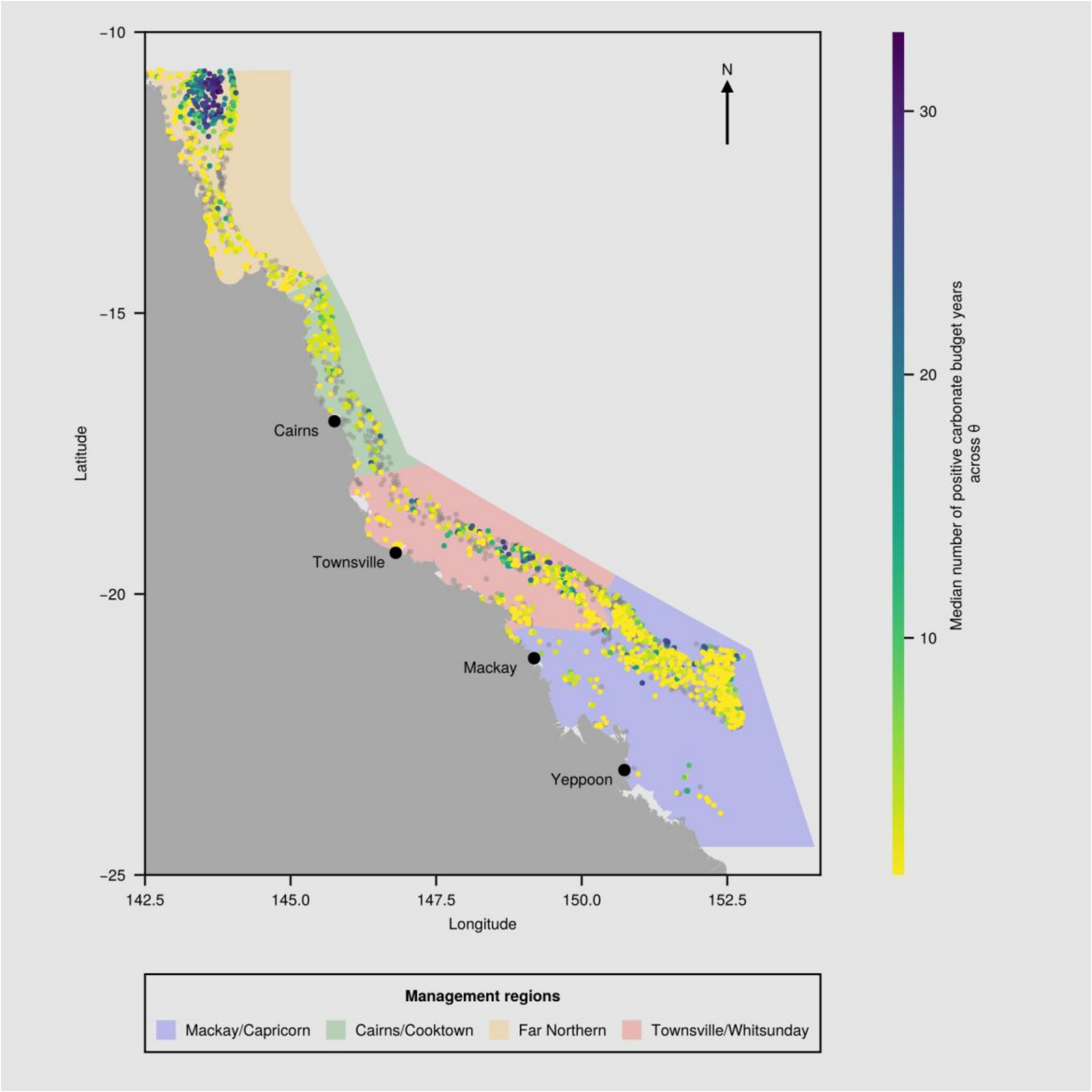


*SFigure 17: Map of the Great Barrier Reef region, including GBRMPA management areas (polygons) and all considered reefs (points), coloured by median number of years exceeding positive carbonate budget threshold. Colour-coded areas outline GBR marine park management regions, where orange represents the Far Northern management region, green represents Cairns/Cooktown, red covers Townsville/Whitsunday and blue covers Mackay/Capricorn (GBRMPA, 2007). Reef centroids are marked individually for visual clarity. Reefs that have a median positive carbonate budget year value of 0 years are coloured in grey. All other reefs are coloured according to their median years across threshold values, with darker colours representing greater number of years with positive carbonate budgets than lighter colours. All 2197 reefs included in this investigation are represented in this figure. This map depicts values from the ACCESS-ESM1-5 GCM. Annotations represent major coastal settlements along the GBR.*

#### Bioregion scale clustering

The following section includes bioregion scale clustering results, full results for other spatial scales (management area and GBR) can be found at Grier and Iwanaga (2025). Results are only shown and discussed for ACCESS-ESM1-5, an intermediate ECS scenario (SFigure 1). Those interested in results for the other assessed GCMs are referred to Grier and Iwanaga (2025). Reefs in the High cluster were consistently found to have the greatest median reef depths across bioregions (SFigure 19). Across bioregions, there were no consistent patterns between reef clusters in mean DHW values (SFigure 20), log10-weighted incoming connectivity (SFigure 22), log10 outgoing connection strength (SFigure 23) and log10 larval retention probability (SFigure 24). Across six of the 29 bioregions reefs in the high cover cluster have lower log10 carrying capacity values than reefs in other clusters (SFigure 21).

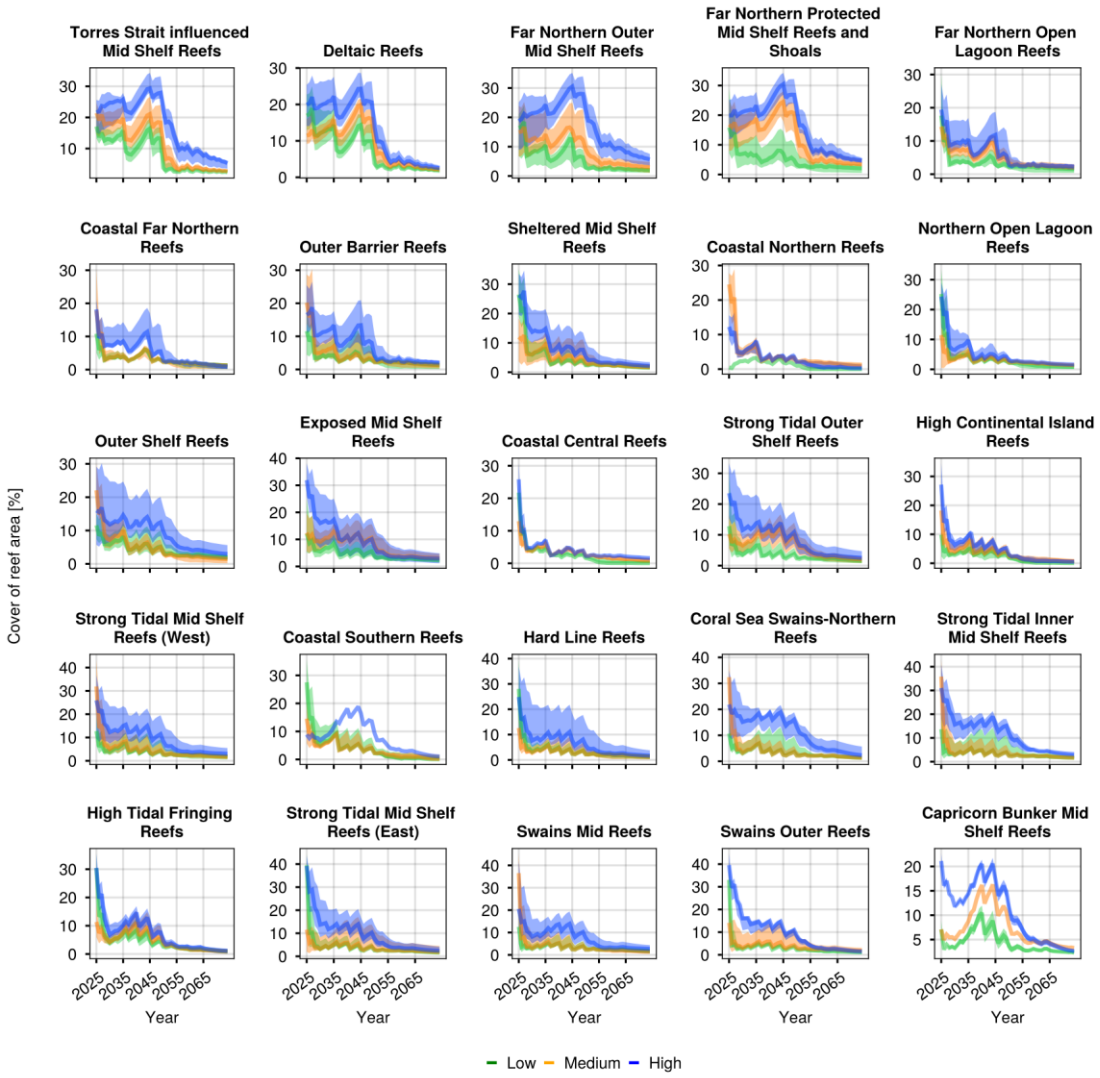


*SFigure 18: Reef area-relative coral cover time series clustered within each bioregion. Time series were clustered based on their behaviour between 2030 and 2060. Time series with comparatively low median area-relative cover is indicated in green, intermediate cover in orange, and high in blue. Solid lines represent the cluster median time series and bands represent the 95th percentile confidence intervals. Displays results for ACCESS-ESM1-5 GCM.*

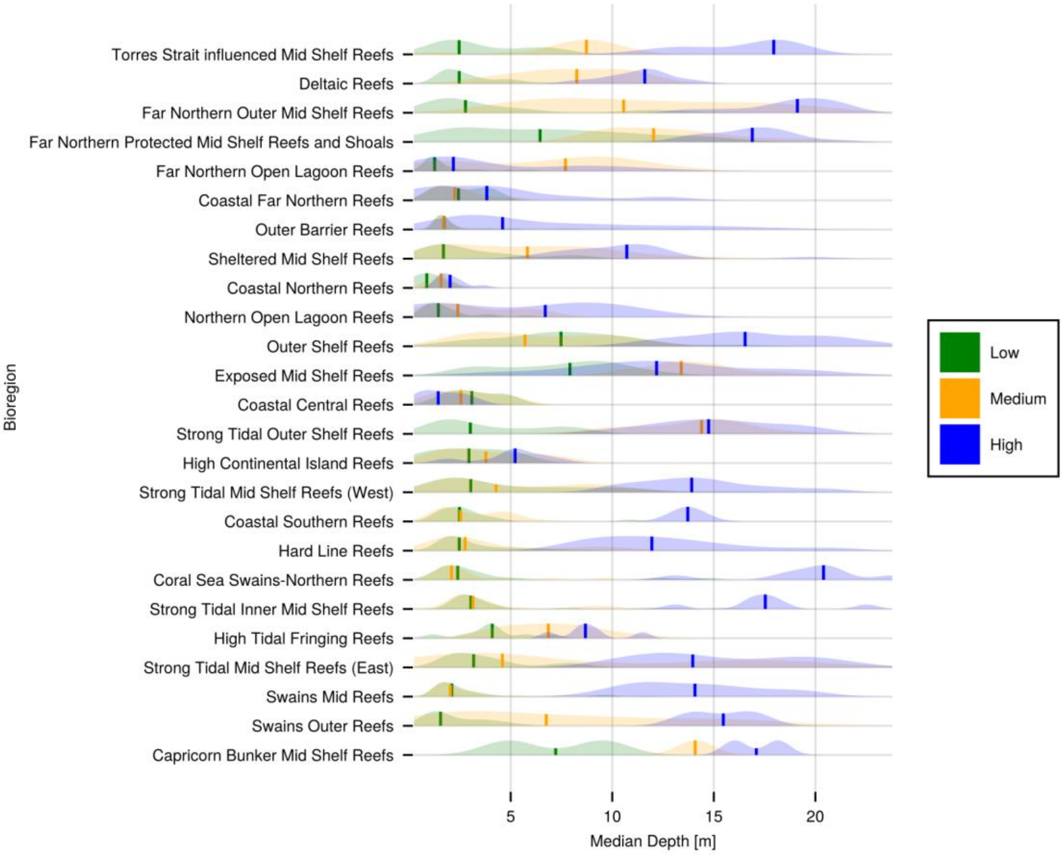


*SFigure 19: Median depth for reefs within each bioregion under ACCESS-ESM1-5, separated into clusters assigned by time series clustering. Time series were clustered based on their behaviour between 2030 and 2060. Time series with comparatively low median area-relative cover is indicated in green, intermediate cover in orange, and high in blue. Solid lines represent the median depth for each cluster across reefs and bands represent the distribution of depth values within each cluster.*

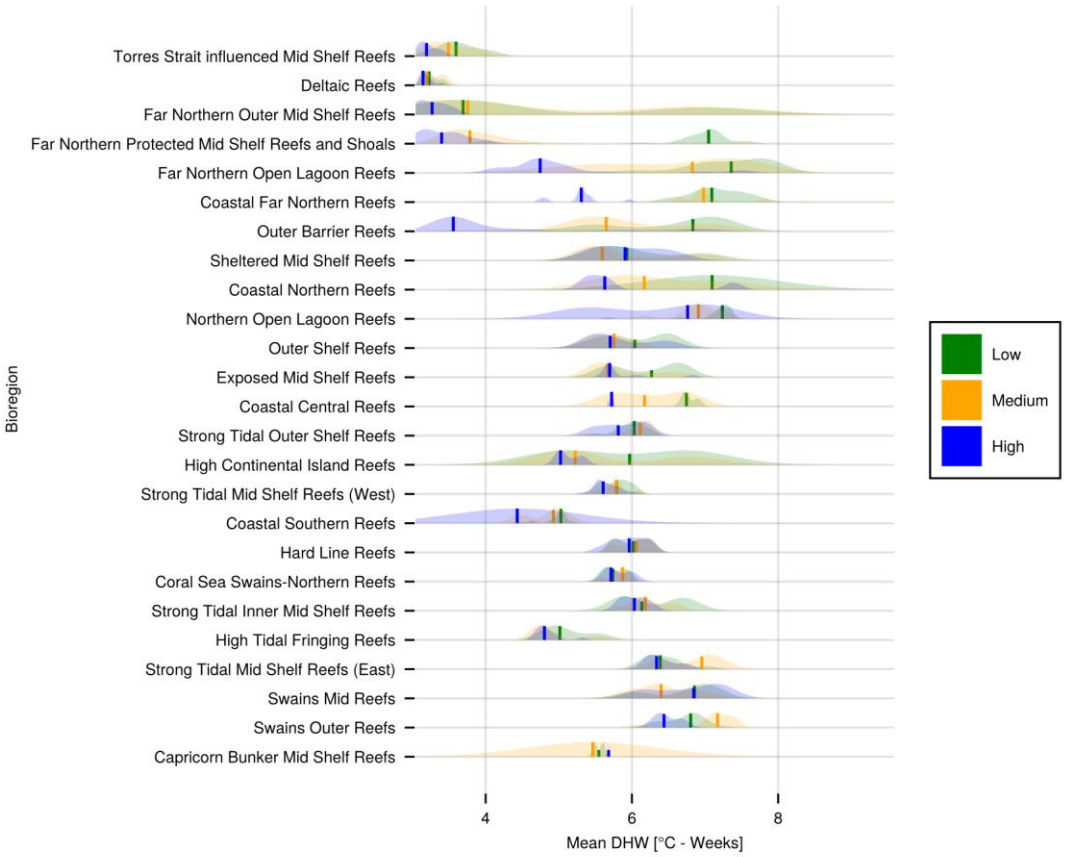


*SFigure 20: Mean time series DHW values for reefs within each bioregion, separated into clusters assigned by time series clustering, under ACCESS-ESM1-5. Time series were clustered based on their behaviour between 2030 and 2060. Time series with comparatively low median area-relative cover is indicated in green, intermediate cover in orange, and high in blue. Solid lines represent the median DHW level for each cluster and bands represent the distribution of mean DHW levels within each cluster.*

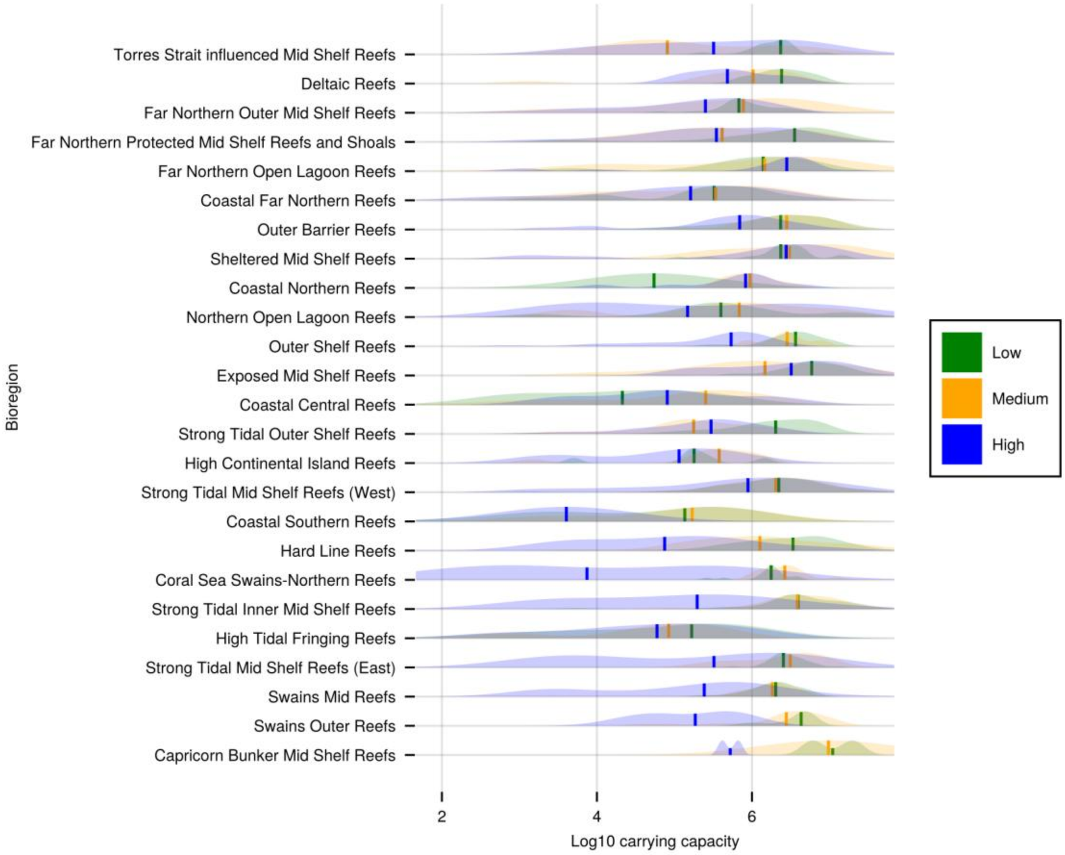


*SFigure 21: Reef carrying capacity (coral habitable area, in base-10 log scale) for reefs within each bioregion, separated into clusters assigned by time series clustering, under ACCESS-ESM1-5. Time series were clustered based on their behaviour between 2030 and 2060. Time series with comparatively low median area-relative cover is indicated in green, intermediate cover in orange, and high in blue. Solid lines represent the median carrying capacity for each cluster and bands represent the distribution of values within each cluster.*

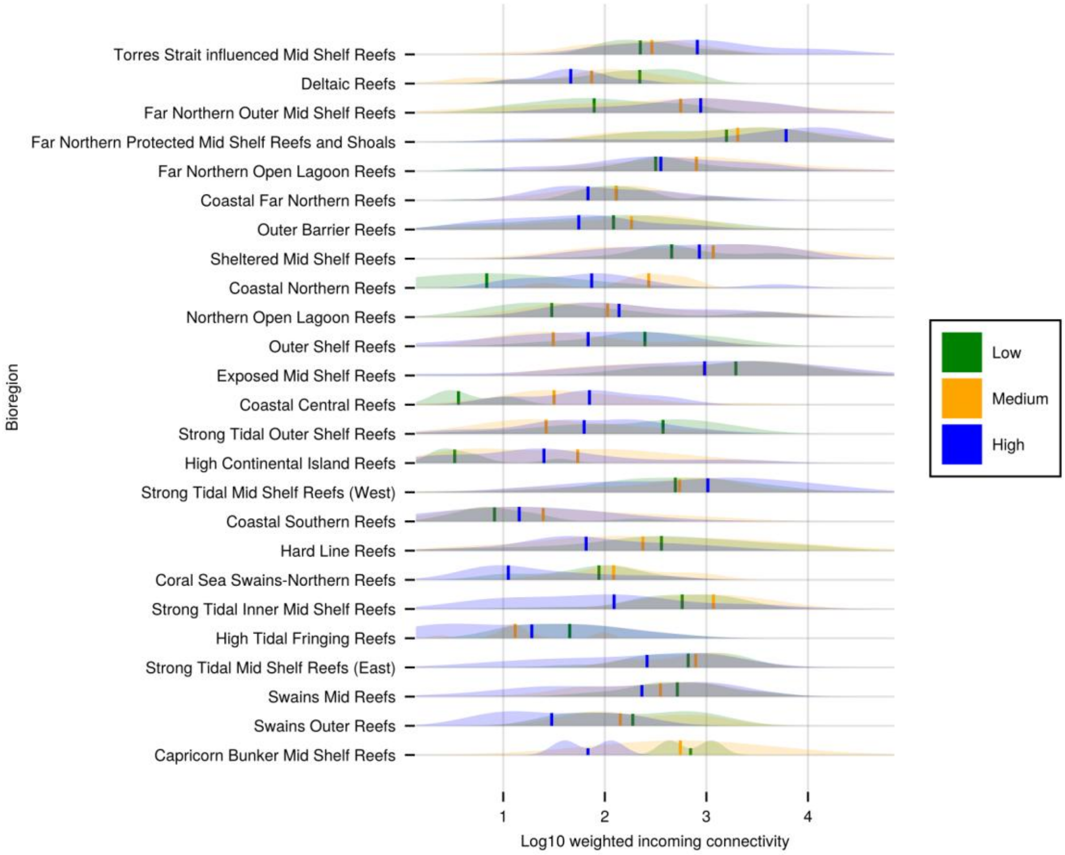


*SFigure 22: Weighted incoming connectivity (in base-10 log scale) for reefs within each bioregion, separated into clusters assigned by time series clustering, under ACCESS-ESM1-5. Time series were clustered based on their behaviour between 2030 and 2060. Time series with comparatively low median area-relative cover is indicated in green, intermediate cover in orange, and high in blue. Solid lines within each violin represent the median base-10 log of weighted incoming connectivity for each cluster and bands represent the distribution of connectivity values within each cluster.*

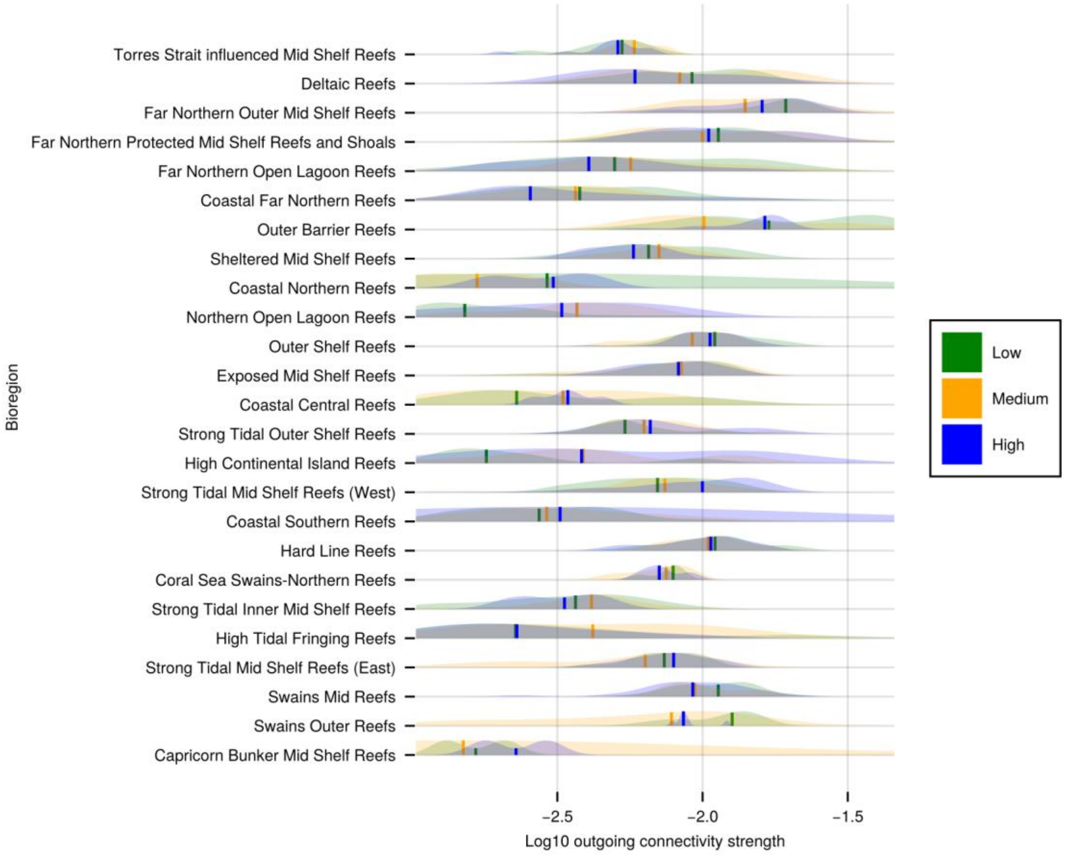


*SFigure 23: Outgoing connection strength (in base-10 log scale) for reefs within each bioregion, separated into clusters assigned by time series clustering, under ACCESS-ESM1-5. Time series were clustered based on their behaviour between 2030 and 2060. Time series with comparatively low median area-relative cover is indicated in green, intermediate cover in orange, and high in blue. Solid lines within each violin represent the median base-10 log of outgoing connection strength for each cluster and bands represent the distribution of connectivity values within each cluster.*

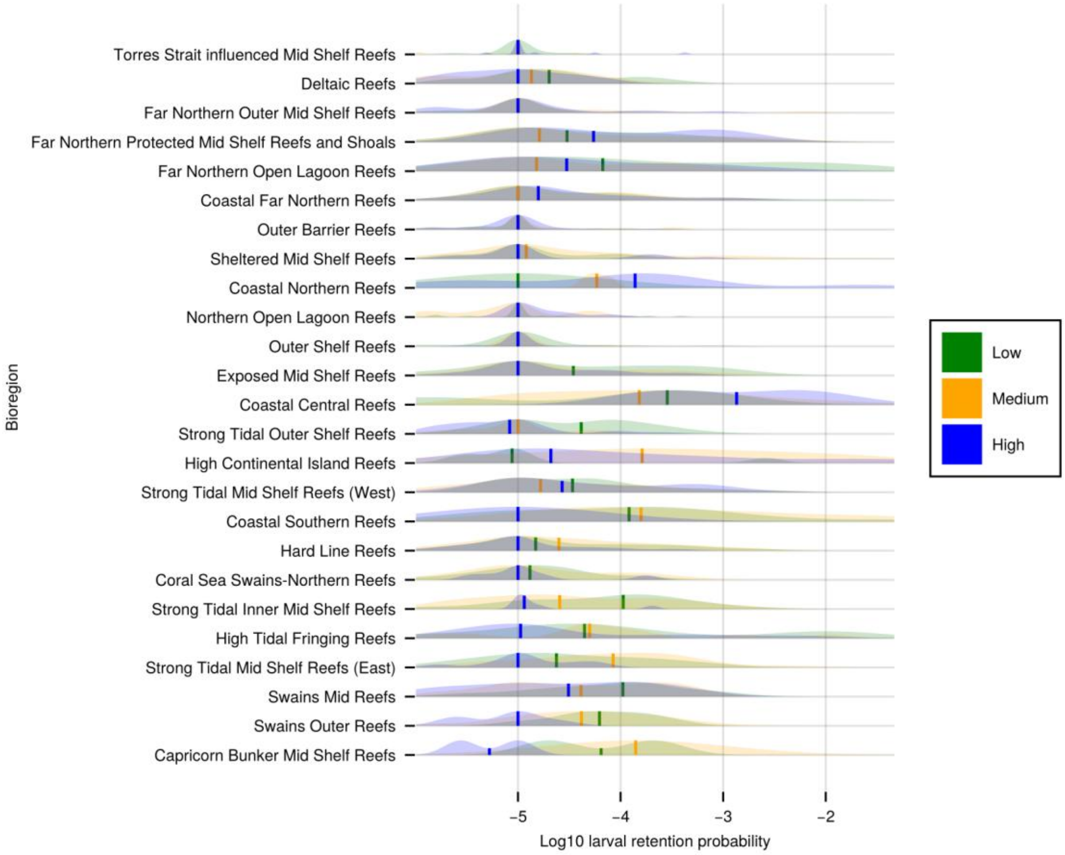


*SFigure 24: Larval retention probability (in base-10 log scale) for reefs within each bioregion, separated into clusters assigned by time series clustering, under ACCESS-ESM1-5. Time series were clustered based on their behaviour between 2030 and 2060. Time series with comparatively low median area-relative cover is indicated in green, intermediate cover in orange, and high in blue. Solid lines within each violin represent the median base-10 log of larval retention probability for each cluster and bands represent the distribution of probability values within each cluster.*

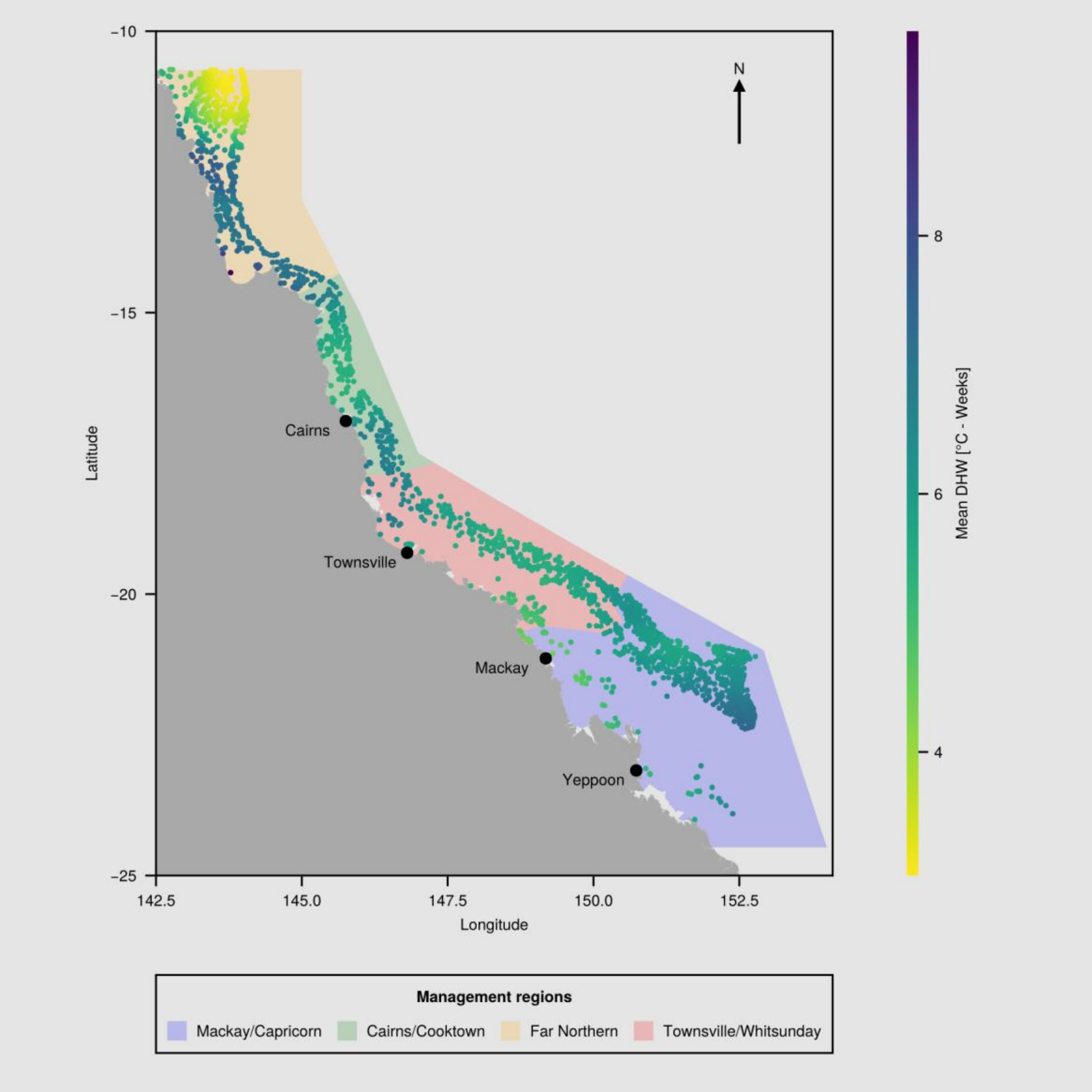


*SFigure 25: Map of the Great Barrier Reef region, including GBRMPA management areas (polygons) and all considered reefs (points), coloured by mean time series DHW value under ACCESS-ESM1-5. Colour-coded areas outline GBR marine park management regions, where orange represents the Far Northern management region, green represents Cairns/Cooktown, red covers Townsville/Whitsunday and blue covers Mackay/Capricorn (GBRMPA, 2007). Reef centroids are marked individually for visual clarity. Darker colours represent higher mean DHW values than lighter colours. All 2197 reefs included in this investigation are represented in this figure. Annotations represent major coastal settlements along the GBR.*

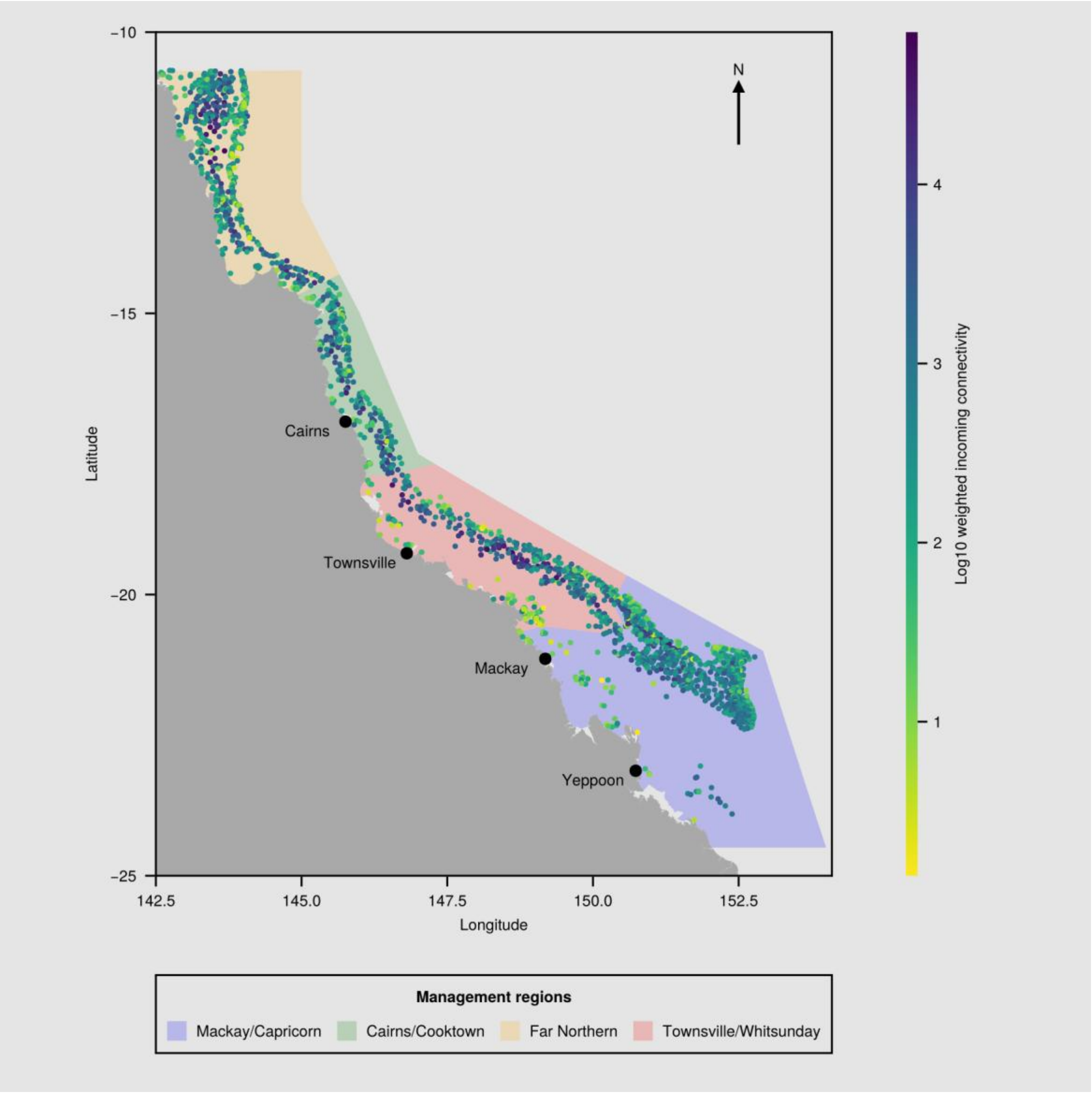


*SFigure 26: Map of the Great Barrier Reef region, including GBRMPA management areas (polygons) and all considered reefs (points), coloured by weighted incoming connectivity levels (Log10) under ACCESS-ESM1-5. Colour-coded areas outline GBR marine park management regions, where orange represents the Far Northern management region, green represents Cairns/Cooktown, red covers Townsville/Whitsunday and blue covers Mackay/Capricorn (GBRMPA, 2007). Reef centroids are marked individually for visual clarity. Darker colours represent higher weighted incoming connectivity values than lighter colours. All 2197 reefs included in this investigation are represented in this figure. Annotations represent major coastal settlements along the GBR.*

#### 142 Random Forest Performance

| | | **Ground Truth** | |
|---|---|---|---|
| **Predicted** | **Low** | **Medium** | **High** |
| **Low** | 1336 | 633 | 138 |
| **Medium** | 297 | 556 | 310 |

| | | | |
|---|---|---|---|
| **High** | 78 | 324 | 722 |

*STable 3: Confusion matrix for the Random Forest classifier used to assess partial dependence. Diagonals indicate correctly classified time series clusters. Columns are the “true” cluster memberships and indicate the samples that were correctly or incorrectly classified as Low/Medium/High. For example, the first column shows the number of samples correctly classified into the Low cluster, and the number of Low samples incorrectly classified as being in either the Medium or High coral cover clusters. There were 4580 samples used for testing.*

| | **Low** | **Medium** | **High** |
|---|---|---|---|
| **Test samples** | 1715 | 1587 | 1278 |
| **Precision** | 0.634 | 0.478 | 0.642 |
| **Recall** | 0.781 | 0.367 | 0.617 |
| **F1** | 0.7 | 0.416 | 0.629 |

*STable 4: Performance metrics for the Random Forest classifier for each of the time series clusters.*
